# Autistic traits influence the strategic diversity of information sampling: insights from two-stage decision models

**DOI:** 10.1101/582783

**Authors:** Haoyang Lu, Li Yi, Hang Zhang

**Author notes:** Corresponding authors: Li Yi, Hang Zhang.

## Abstract

Information sampling can reduce uncertainty in future decisions but is often costly. To maximize reward, people need to balance sampling cost and information gain. Here we aimed to understand how autistic traits influence the optimality of information sampling and to identify the particularly affected cognitive processes. Healthy human adults with different levels of autistic traits performed a probabilistic inference task, where they could sequentially sample information to increase their likelihood of correct inference and may choose to stop at any moment. We manipulated the cost and evidence associated with each sample and compared participants’ performance to strategies that maximize expected gain. We found that participants were overall close to optimal but also showed autistic-trait-related differences. Participants with higher autistic traits had a higher efficiency of winning rewards when the sampling cost was zero but a lower efficiency when the cost was high and the evidence was more ambiguous.

Computational modeling of participants’ sampling choices and decision times revealed a two-stage decision process, with the second stage being an optional second thought. Participants may consider cost in the first stage and evidence in the second stage, or in the reverse order. The probability of choosing stopping at a specific stage increases with increasing cost or increasing evidence. Surprisingly, autistic traits did not influence the decision in either stage. However, participants with higher autistic traits inclined to consider cost first, while those with lower autistic traits considered cost or evidence first in a more balanced way. This would lead to the observed autistic-trait-related advantages or disadvantages in sampling optimality, depending on whether the optimal sampling strategy is determined only by cost or jointly by cost and evidence.

**Author Summary:** Children with autism can spend hours practicing lining up toys or learning all about cars or lighthouses. This kind of behaviors, we think, may reflect suboptimal information sampling strategies, that is, a failure to balance the gain of information with the cost (time, energy, or money) of information sampling. We hypothesized that suboptimal information sampling is a general characteristic of people with autism or high level of autistic traits. In our experiment, we tested how participants may adjust their sampling strategies with the change of sampling cost and information gain in the environment. Though all participants were healthy young adults who had similar IQs, higher autistic traits were associated with higher or lower efficiency of winning rewards under different conditions. Counterintuitively, participants with different levels of autistic traits did not differ in the general tendency of oversampling or undersampling, or in the decision they would reach when a specific set of sampling cost or information gain was considered. Instead, participants with higher autistic traits consistently considered sampling cost first and only weighed information gain during a second thought, while those with lower autistic traits had more diverse sampling strategies that consequently better balanced sampling cost and information gain.

## Introduction

Information helps to reduce uncertainty in decision making but is often costly to collect. For example, to confirm whether a specific tumor is benign or malignant may require highly invasive surgery procedures. In such cases, it can be more beneficial to tolerate some degree of uncertainty and take actions first. To maximize survival, humans and animals need to balance the cost and benefit of information sampling and sample the environment optimally [1,2].

However, autism spectrum disorder (ASD)—a neurodevelopmental disorder characterized by social impairments and repetitive behaviors [3]—seem to be accompanied by suboptimal information sampling, though in various guises. For example, individuals with repetitive behaviors tend to spend time on redundant information that helps little to reduce uncertainty [4]. Eye-tracking studies reveal that people with ASD have atypical gaze patterns in ambiguous or social scenes, that is, they sample the visual environment in an inefficient way [5,6]. According to the recently developed Bayesian theories of ASD that explain a variety of perceptual, motor, and cognitive symptoms [7–13], deviation from Bayesian optimality in information processing is primary to ASD [4,14–17]. In this Bayesian framework, information sampling is referred as “disambiguatory active inference” [4] and plays an important role in guiding the subsequent inferences or decisions. We hereby conjectured that ASD symptoms such as repetitive behaviors and ineffecient gaze patterns reflect general impairments in information sampling.

The autistic traits of the whole population form a continuum, with ASD diagnosis usually situated on the high end [18–24]. Moreover, autistic traits share genetic and biological etiology with ASD [25]. Thus, quantifying autistic-trait-related differences in healthy people can provide unique perspectives as well as a useful surrogate for understanding the symptoms of ASD [23,26].

The present study is aimed to understand how autistic traits in typical people may influence their optimality of information sampling. In particular, we focused on the situation where information can be used to improve future decisions (e.g. [27–29], in contrast to non-instrumental information gathering such as [30–39]) and hypothesized that individuals with high autistic traits may deviate more from optimality in information sampling.

Possible suboptimality may arise from a failure of evaluating sampling cost or information gain, or improper trading off the two, or a greater noise [27]. To investigate these possibilities, we tested healthy adults of different levels of autistic traits in an information sampling task adapted from [40,41]: On each trial of the experiment, participants could draw samples sequentially to accumulate evidence for a probabilistic inference and would receive monetary rewards for correct inferences. Each additional sample may increase their probability of correct inference but also incur a fixed monetary cost. In order to maximize expected gain, participants should draw fewer samples when each sample had higher cost or provided higher evidence, and vice versa. We manipulated the cost and evidence per sample and compared participants’ performance to optimality. We found that different levels of autistic traits were accompanied by different extents of deviation from optimality. Compared to their peers, participants with higher level of autistic traits received higher rewards in the zero-cost conditions due to less undersampling, where the optimal strategy was to sample as many as possible, but meanwhile lower rewards in the high-cost, low-evidence condition due to more oversampling, where the optimal strategy would sacrifice accuracy to save cost.

What cognitive processes in information sampling are particularly affected by autistic traits? Through computational modeling, we further decomposed participants’ sampling choices into multiple sub-processes and found that the influence of autistic traits was surprisingly selective and subtle. In particular, participants’ sampling choices could be well described by a two-stage decision process: When the first decision stage does not reach the choice of stopping sampling, a second decision stage is probabilistically involved to arbitrate, which offers a second chance to consider stopping sampling. The two stages were independently controlled by cost and evidence and neither stage showed autistic-trait-related differences. What varied with levels of autistic traits was the strategic diversity: Participants with higher autistic traits were more likely to always consider cost in the first stage and evidence in the second, while those with lower autistic traits had a larger chance to use the reverse order as well. As a consequence, the former would perform better when the optimal strategy does not depend on evidence, while the latter would do better when the optimal strategy is determined jointly by cost and evidence.

## Results

One hundred and four healthy young adults participated in our experiment, whose autistic traits were measured by the self-reported Autism Spectrum Quotient (AQ) questionnaire [18]. The computerized experimental task is illustrated in Fig 1a. On each trial, participants first saw two jars filled with opposite ratios of pink and blue beads and were told that one jar had been secretly selected by the experimenter. They could sample up to 20 beads sequentially with replacement from the selected jar to infer which jar had been selected. Each key press would randomly sample one bead and participants could decide to stop sampling at any moment. For each correct inference, participants would receive 10 points minus the total sampling cost. Their goal was to earn as many points as possible, which would be redeemed into monetary bonus in the end. The cost of sampling one bead could be 0, 0.1, or 0.4 points, referred below as zero-, low-, and high-cost conditions respectively. The pink-to-blue ratios of the two jars could be 60%:40% vs. 40%:60%, or 80%:20% vs. 20%:80%, which corresponded to lower (60/40) or higher (80/20) evidence per sample favoring one jar against another. The sample size that maximizes expected gain would change with the cost and evidence conditions (Fig 1b, see Methods).

**Fig 1.**
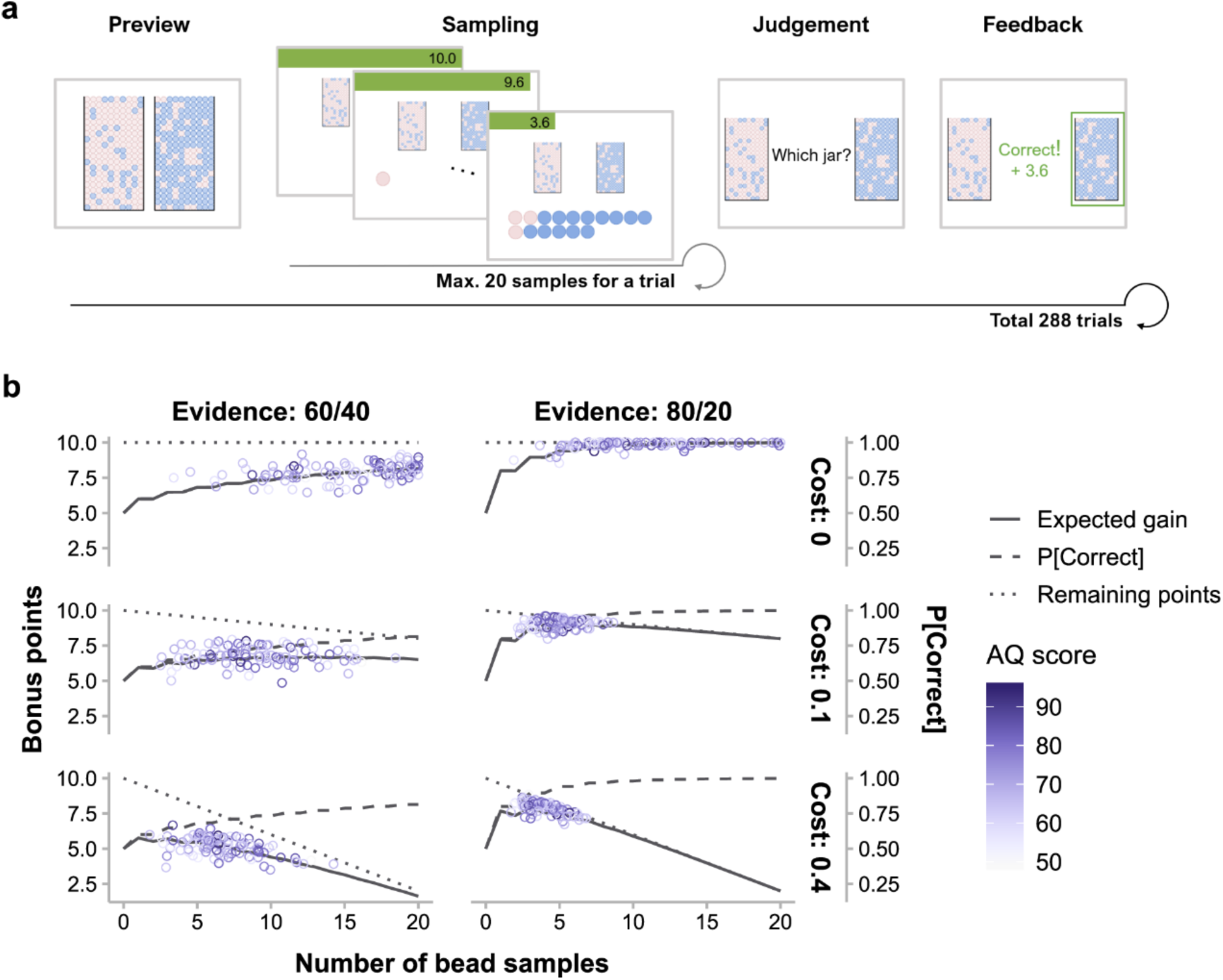
The bead-sampling task. (a) Time course of one trial. “Preview” informed the participant of the pink-to-blue ratios of the two jars (80%:20% vs. 20%:80% in this example, corresponding to the high-evidence condition). Then the participant could sample beads from the unknown pre-selected jar one at a time up to 20 beads (“sampling”) or quit sampling at any time. Afterward, the participant judged which jar had been selected (“judgment”). Feedback followed, showing the correctness of judgment and winning of the current trial. Feedback was presented for 1 s, whereas preview, sampling, and judgment were self-paced. During sampling, the remaining bonus points (green bar), as well as the array of bead samples, were visualized and updated after each additional sample. (b) Optimal sampling strategy vs. participants’ performance for each of the six cost-by-evidence conditions. On a specific trial, the expected probability of correctness (dashed lines) and the remaining bonus points (dotted lines) are respectively increasing and decreasing functions of the number of bead samples. The expected gain (solid lines), as their multiplication product, first increases and then decreases with the number of samples. Note that the sample size that maximizes expected gain varies across different cost and evidence conditions. Each circle represents a participant with the color indicating their AQ score.

### Sampling optimality may increase or decrease with autistic traits in different conditions

We computed efficiency—the expected gain for participants’ sample sizes divided by the maximum expected gain—to quantify the optimality of participants’ sampling choices and used linear mixed model analyses to identify the effects of AQ and its interactions with sampling cost and information gain (LMM1 for efficiency, see Methods). Participants’ efficiency (Fig 2a) was on average 94% (i.e. close to optimality) but decreased with increasing cost (*F*_2,100.98_ = 65.38, *p* < .001) or decreasing evidence (*F*_1,101.88_ = 124.95, *p* < .001), and decreased more dramatically when high cost and low evidence co-occurred (interaction *F*_2,202.89_ = 123.20, *p* < .001). Though participants with different AQ did not differ in overall efficiency, AQ influenced efficiency through its interaction with cost and evidence (three-way interaction *F*_2,203.45_ = 5.60, *p* = .004). As post hoc comparisons, we compared the regression slope of AQ—the change in efficiency with one unit of increase in AQ—across conditions (Fig 2d). Under the low-evidence conditions, the slope was more negative under high cost than under zero (*t*_137.82_ = −3.16, *p* = .005) or low cost (*t*_151.58_ = −2.64, *p* = .023). No significant differences were found among different costs in the high evidence conditions. In almost all conditions the slope was non-negative or even significantly positive (i.e. the zero cost, low evidence condition, *t*_136.08_ = 2.11, *p* = .037), indicating higher efficiency for participants with higher AQ. However, when sampling was both costly and little informative (i.e. the high-cost, low-evidence condition), the efficiency decreased with AQ (*t*_121.32_ = −2.51, *p* = .014). We verified these AQ-related differences in an alternative analysis, where we divided participants evenly into three groups of low, middle, and high AQ scores and found similar results (S1 Fig).

**Fig 2.**
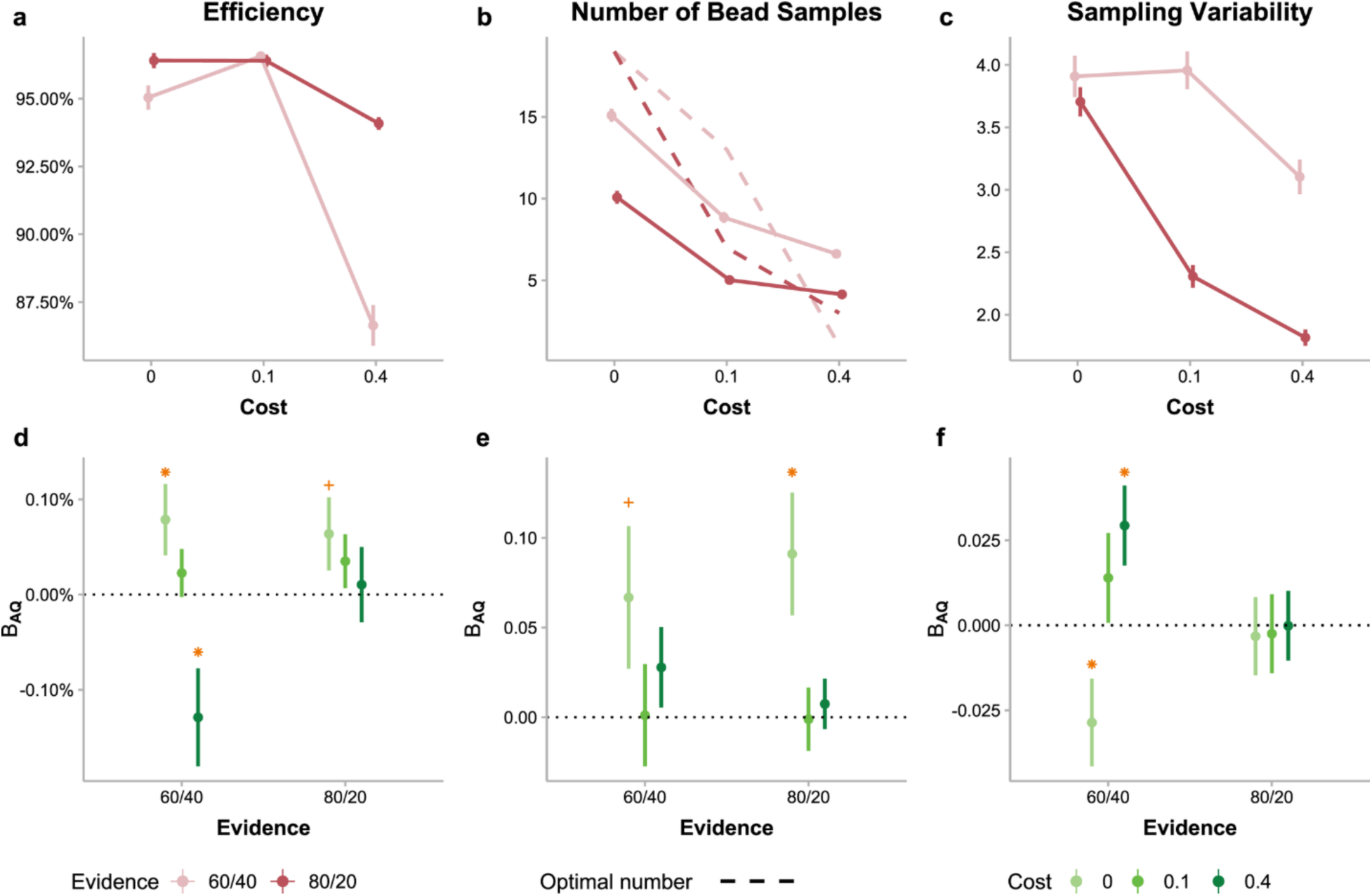
Optimality of sampling performance and the effects of autistic traits. (a) Sampling efficiency varied with cost (abscissa) and evidence (different colors) conditions. Participants’ efficiency was on average 94% (i.e. close to optimality) but decreased with increasing cost or decreasing evidence, and decreased more dramatically when high cost and low evidence co-occurred. (b) The mean number of bead samples participants drew in a condition (solid lines) decreased with increasing cost or increasing evidence. Compared to the optimal number of samples (dashed lines), participants undersampled in the zero- or low-cost conditions while oversampled in the high-cost conditions. (c) Sampling variability (standard deviation of the numbers of samples drawn across trials) varied with cost and evidence conditions. Error bars in (a) – (c) denote between-subject standard errors. (d) – (f) Effects of AQ levels on participants’ sampling performance in different cost (different colors) and evidence (abscissa) conditions. Β_AQ_ is the unstandardized coefficient of AQ indicating how much the efficiency (d), number of samples (e), and sampling variability (f) would change when AQ increases by one unit. Error bars represent standard errors of the coefficients. Orange asterisk: *p* < .05, orange plus: *p* < .1.

The overall high efficiency was accompanied by adaptive sampling behaviors that were modulated by both sampling cost and information gain: Participants drew fewer samples in costlier or more informative conditions as the optimal strategy would require (Fig 2b). We quantified participants’ sampling behaviors in a particular condition using two measures: sampling bias (the actual number of sampling minus the optimal number of sampling, denoted 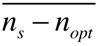) and sampling variability (standard deviation of the actual numbers of sampling, denoted *SD*(*n_s_*)).

A linear mixed model analysis on 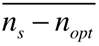 (LMM2, see Methods) showed main effects of cost (*F*_2,100.93_ = 752.65, *p* < .001) and evidence (*F*_1,101.98_ = 177.48, *p* < .001), as well as their interactions (*F*_2,202.97_ = 546.59, *p* < .001). Similar to its influence on efficiency, AQ did not lead to a general tendency of more oversampling or undersampling but had significant interactions with cost (*F*_2,101.13_ = 3.99, *p* = .022). In particular, the slope of AQ for 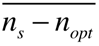 (Fig 2e) was more positive for the zero-cost than for the low-cost condition (*t*_101.90_ = 2.61, *p* = .025). Under zero cost, given that participants tended to undersample (Fig 2b), a positive slope of AQ (*t*_110.02_ = 2.67, *p* = .009 for high evidence and *t*_107.86_ = 1.68, *p* = .096 for low evidence) implies less undersampling for participants with higher AQ.

According to a similar linear mixed model analysis on *SD*(*n_s_*) (LMM3, see Methods), the main effects of cost (*F*_2,100.86_ = 57.13, *p* < .001) and evidence (*F*_1,101.89_ = 161.78, *p* < .001) as well as their interactions (*F*_2,203.43_ = 33.51, *p* < .001) were significant (Fig 2c). Again, AQ influenced sampling variability through its interaction with cost and evidence (three-way interaction *F*_2,204.09_ = 5.27, *p* = .006). Post hoc comparisons showed that the slope of AQ for sampling variability was more negative under zero cost than under low (*t*_172.54_ = −2.43, *p* = .042) or high cost (*t*_188.90_ = −3.51, *p* = .002) in the low-evidence conditions but was little influenced by cost in the high-evidence conditions (Fig 2f). In the low-evidence conditions, the observed slopes imply that higher AQ led to lower sampling variability under zero cost (*t*_140.52_ = −2.22, *p* = .028) but higher sampling variability under high cost (*t*_154.23_ = 2.50, *p* = .014).

Taken together, participants with different levels of AQ differed in both the mean and SD of sample sizes. Participants with higher AQ had higher efficiency in the zero-cost, low-evidence condition, which was associated with less undersampling and lower sampling variability. Meanwhile, higher AQ corresponded to lower efficiency and higher sampling variability in the high-cost, low-evidence condition.

### Bimodal decision times suggest two consecutive decision processes

Decision time (DT) for a specific sample—the interval between the onset of last bead sample (or, for the first sample, the start of the sampling phase) and the key press to draw the sample—provided further information about the cognitive process underlying sampling choices. Though decision or response times usually have a positively skewed unimodal distribution and are close to Gaussian when log-transformed [42,43], the log-transformed DTs for continuing sampling in our experiment had a bimodal distribution (Hartigan’s dip test for multimodality, *D* = 0.004, *p* < .001), well fitted by a mixture of two Gaussian distributions (Fig 3a). Such bimodality was evident in the low-cost and high-cost conditions (low-cost, low-evidence: *D* = 0.013, *p* < .001; low-cost, high-evidence: *D* = 0.009, *p* < .001; high-cost, low-evidence: *D* = 0.014, *p* < .001; high-cost, high-evidence: *D* = 0.015, *p* < .001), but was barely palpable in the zero-cost conditions (zero-cost, low-evidence: *D* = 0.002, *p* = .11; zero-cost, high-evidence: *D* = 0.001, *p* = .95), where the first peak was dominant. Similar bimodal distributions were observed for individual participants (S2 Fig) and could not simply be artifacts of data aggregation.

**Fig 3.**
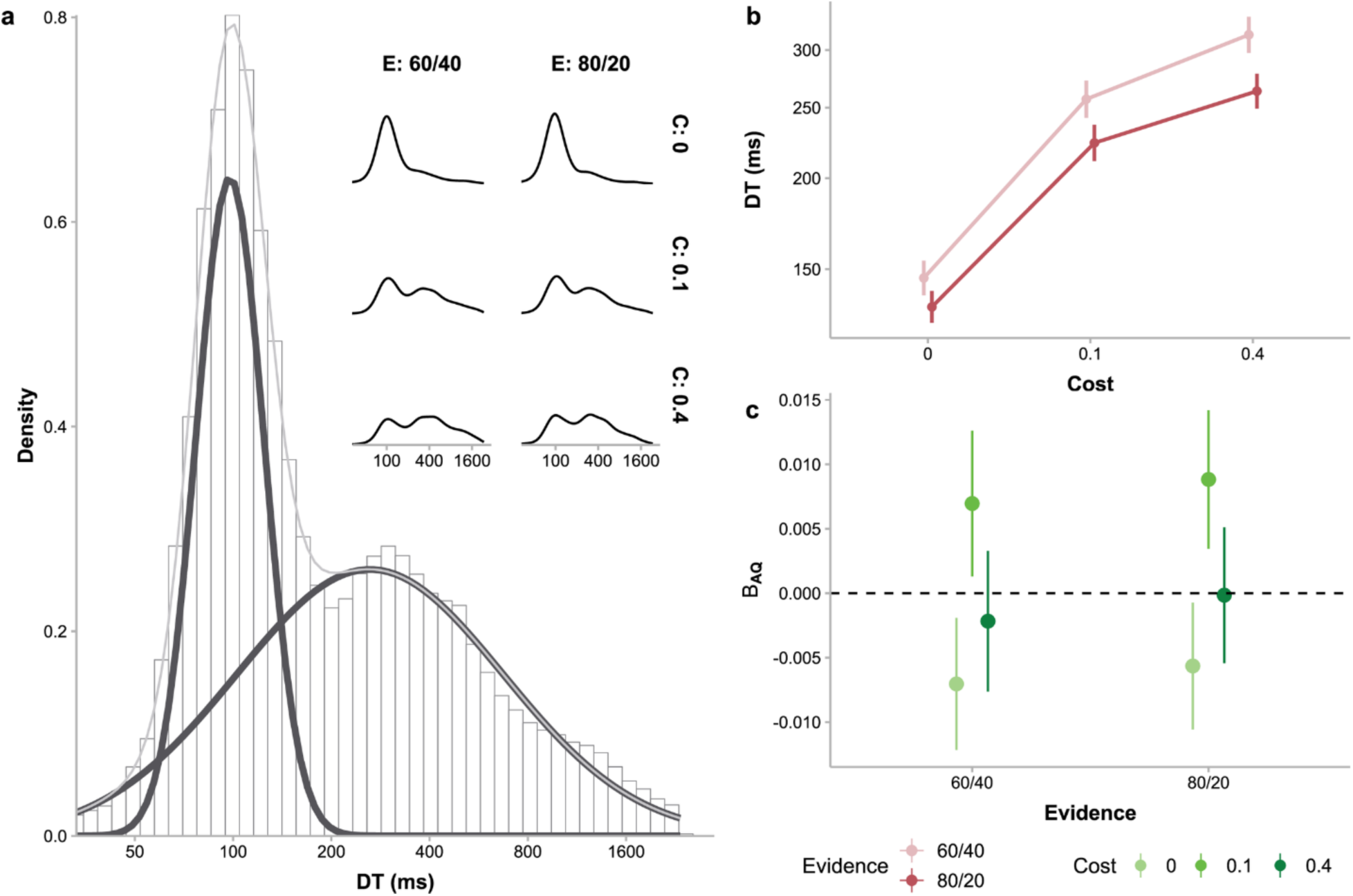
Decision time (DT) for each sampling. (a) The distributions of DTs aggregated over all participants (main plot) and for each cost and evidence condition (insets). In the main plot, the distribution of DTs (histogram) was clearly bimodal, well fitted by a Gaussian mixture (gray curve) with two Gaussian components (black curves). Such bimodality was also visible in most inset plots, though the relative weights of the two components varied with experiment conditions. (b) Mean DTs varied with cost (abscissa) and evidence (different colors) conditions. Error bars represent between-subject standard errors. (c) Effects of AQ levels on participants’ DTs in different cost (different colors) and evidence (abscissa) conditions. Β_AQ_ is the unstandardized coefficient of AQ indicating how much the mean DT in a condition would change when AQ increases by one unit. Error bars represent standard errors of the coefficients.

Linear mixed model analysis (LMM4) showed that the mean DTs (Fig 3b) increased with cost (*F*_2,101_ = 120.62, *p* < .001) and decreased with evidence (*F*_1,102_ = 165.85, *p* < .001). The difference between different evidence conditions was also larger for higher sampling cost (interaction *F*_2,204_ = 14.65, *p* < .001). Moreover, there was a significant interaction between cost and AQ (*F*_2,101_ = 6.22, *p* = .003): DTs tended to decrease with AQ under zero cost but increase with AQ under low cost (Fig 3c, slope difference between these two conditions reached significance, *t*_102_ = 3.45, *p* = .002).

The DTs within the same trial changed with sample number (LMM5, *F*_19,10805525.21_ = 24.5, *p* < .001). Post hoc contrasts showed significantly negative linear trends (S3 Fig, *t*_4323568_ = −12.26, *p* < .001), indicating that sampling decisions in a trial became faster after more samples were drawn. AQ significantly moderated the effect of sample number (interaction *F*_19,1498809.98_ = 1.66, *p* = .035), with higher AQ associated with a flatter trend (*t*_4628456_ = 3.62, *p* = .002). In other words, participants with higher AQ tended not to speed up their decisions as much as those with lower AQ.

A straightforward explanation for the bimodal DT distribution would be a probabilistic mixture of two cognitive processes. Next, we used computational modeling to explore the possibility of two decision stages and showed that it could quantitatively predict the effects of cost and evidence as well as the bimodal distribution of DTs.

### Sampling is controlled by cost and evidence in two separate stages

We considered a variety of models for sampling choices, which fell into two categories: one-stage models and two-stage models (Fig 4a, see Methods). In one-stage models, the choice of whether to take a *(j+1)*-th sample after *j* samples is modeled as a Bernoulli random variable, with the probability of stopping controlled by cost- and evidence-related factors, including the expected cost and evidence for the prospective sample and the total cost and evidence of existing samples. To separate the influences of different factors on participants’ sampling choices, we constructed a set of one-stage models that are controlled either by cost-related factors, or by evidence-related factors, or by both. To test the possibility that people of higher autistic traits may overweight recent evidence in evidence integration [4,17], we also considered models with an evidence decay parameter, in which the weight for an earlier sample decays as a function of the number of samples thereafter.

**Fig 4.**
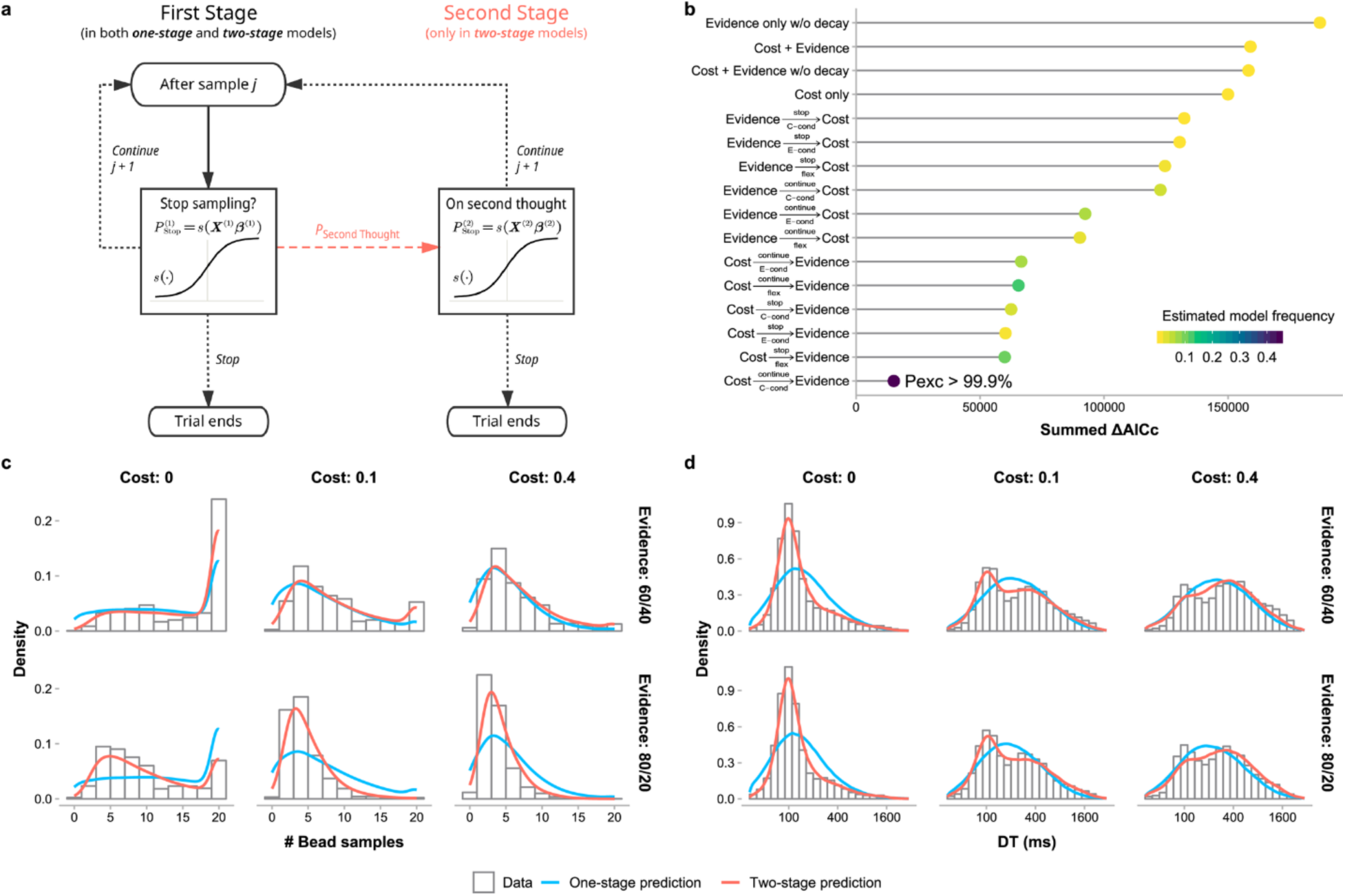
Computational modeling of sampling choices and decision times. Schematic of one-stage and two-stage models. One-stage models only consist of the steps on the left-hand side: Each time a participant decides whether to stop or continue sampling, the probability of stopping is a sigmoid function of a linear combination of multiple decision variables. Two-stage models assume that participants may probabilistically have a second thought to reconsider the choice (the coral dashed arrow). The second stage (on the right-hand side) works in the same way as the first stage but the two stages are controlled by different sets of decision variables. (b) Results of model comparison based on the joint fitting of choice and DT. The ΔAICc for a specific model was calculated for each participant with respect to the participant’s best-fitting model (i.e. lowest-AICc) and then summed across participants. Both fixed-effects (summed ΔAICc: lower is better) and random-effects (estimated model frequency: higher is better) comparisons revealed that the best-fitting model was a two-stage model with cost-related variables considered in the first stage and evidence-related variables in the second stage (i.e. 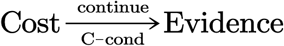). The best one-stage model was the model involving only cost-related decision variables (i.e. Cost only). See Methods (or S1 Table) for the description of each model. Estimated model frequency (color coded) is a random effects measure of the proportion of participants best fit by the model. (c) Distribution of sample sizes (i.e. number of bead samples) for each condition: data vs. model predictions. (d) Distribution of DTs for each condition: data vs. model predictions. The best-fitted two-stage model (red curves) well predicted the observed distributions (histograms) of sample sizes and DTs for each cost and evidence condition, including the bimodality of the observed DT distributions, while the best-fitted one-stage model (blue curves) failed to do so. Both data and model predictions were aggregated across participants.

In two-stage models of sampling choices, we assumed that deciding whether to stop or continue sampling may involve two consecutive decision stages, where the decision in the first stage can either be final or be re-evaluated in an optional second stage. Whether to enter the second stage is probabilistic, conditional on the decision reached in the first stage. The decisions in the two stages are independent and controlled separately by the cost- and evidence-related factors and are subject to evidence decay. In other words, the decision in each stage is similar to that of a one-stage model. We considered 12 different two-stage models whose assumptions differ in three dimensions (see Methods): (1) which factors control the first stage and which control the second stage (cost-first or evidence-first), (2) what kind of decision in the first stage (continuing or stopping sampling) has a chance to trigger the second stage, and what determines the probability to enter the second stage (“second-thought probability”) after a qualified first-stage decision. For example, the best-fitting second-stage model described below, denoted 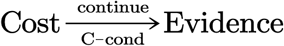, has the following assumptions: cost-related factors control the first stage and evidence-related factors control the second stage. If stopping sampling is the decision in the first stage, it is finalized and there is no second stage; otherwise, either continuing sampling becomes the final decision, or the decision is re-evaluated in the second stage, with the second-thought probability determined by the cost condition (i.e. three different second-thought probabilities for the zero-, low-, and high-cost conditions).

We fit all the models to participants’ sampling choices separately for each participant using maximum likelihood estimates. For each fitted choice model, with some additional assumptions, we were able to model participants’ DTs and fit the additional DT parameters using maximum likelihood estimates as well (see Methods). The sum of the log likelihoods for choices and DTs was used for further model comparisons, which was mathematically equivalent to the log likelihood from modeling the joint distribution of choices and RTs (see Methods for proof). We compared the models in goodness-of-fit using the Akaike Information Criterion corrected for small samples (AICc) [44,45]. The ΔAICc for a specific model was calculated for each participant with respect to the participant’s best-fitting model (i.e. lowest-AICc) and then summed across participants. We also used the group-level Bayesian model selection [46,47] for random effects model comparisons and plot each model’s estimated model frequency—a random effects measure of the proportion of participants best fit by the model. Among the four one-stage models (Fig 4b), the best model (i.e. model with the lowest summed ΔAICc) was the one that is influenced by cost only (denoted *Cost only*). However, the two-stage models, all of which were controlled by the same cost- and evidence-related factors as the one-stage models, fit much better to participants’ choices and DTs than the best one-stage model. The best two-stage model was 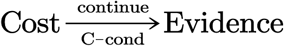 (described above), which best accounted for 50% of the 104 participants (estimated model frequency = 44.6%) and whose probability of outperforming all the other 15 models (protected exceedance probability) approached 1. Model comparisons based on the Bayesian Information Criterion (BIC) [48,49] led to similar results (see S4 Fig for group and individual participants’ ΔAICc and ΔBIC).

When two-stage models were fit to participants’ DTs, the second-thought probabilities were estimated exclusively from choices and not free parameters adjustable by DTs (see Methods). However, predictions of the 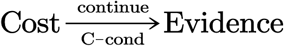 model agreed well not only with participants’ choices but also with their bimodal DTs (Fig 4cd, see S2 & S5 Fig for individual plots) and the decrease of DT with sample number (S3 Fig). This further supports our hypothesis that the observed bimodal DT distribution arises from a two-stage decision process.

As additional evidence for the link between two-stage decisions and bimodal RTs, the mean DT—as a proxy for the proportion of slow decisions—increased with the probability of using the second stage (Fig 5; *r*_s_ = .60, *p* < .001). The positive correlation also held for each separate cost condition (zero cost: *r_s_* = .44, *p* < .001; low cost: *r_s_* = .35, *p* < .001; high cost: *r_s_* = .22, *p* = .027). Moreover, the effects of cost on mean DT (LMM4, as we reported earlier) could be partly explained away by the effect of second-thought probability when the latter was added as a predictor (LMM6; second-thought probability and its interaction with evidence, *F*_1,78.06_ = 47.74, *p* < .001 and *F*_1,284.99_ = 25.76, *p* < .001 respectively; cost and its interaction with evidence, *F*_2,73.75_ = 2.43, *p* = .09 and *F*_2,233.83_ = 2.59, *p* = .08).

**Fig 5.**
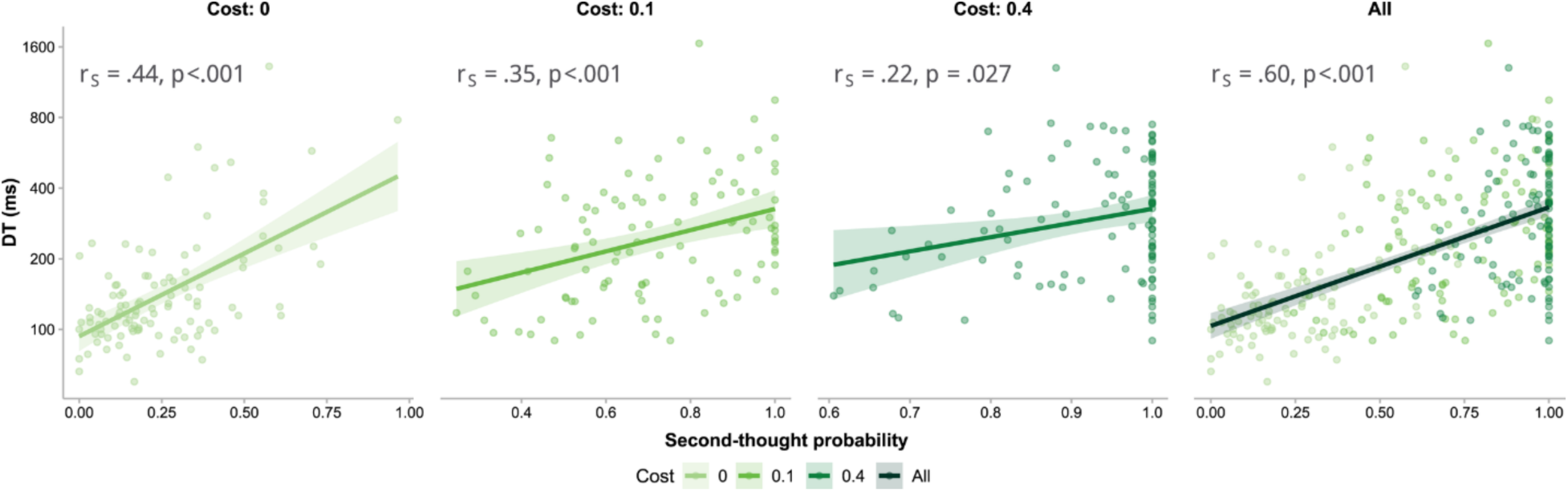
Positive correlations between mean decision time and second-thought probability. According to two-stage models, mean DT—as a proxy for the proportion of slow decisions— should increase with the probability of using the second stage. Indeed, mean DT and second-thought probability were positively correlated, separately for each cost condition (the first three panels) and when aggregated across all cost conditions (the last panel), thus providing additional support for the two-stage decision process. Each dot is for one participant in one specific cost condition. Lines and shaded areas respectively represent regression lines and standard errors. The *r*refers to Spearman’s correlation coefficient.

### Autistic traits influence the strategic diversity of sampling decisions

What individual differences in the decision process may relate to the autistic-trait-related effects on the optimality of sampling choices? We first examined the estimated parameters of the best model (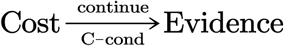), which allowed us to characterize individual participants’ sampling choices from three aspects: cost- or evidence-related weights (11 parameters), second-thought probabilities (three parameters separately for the three cost conditions), and evidence decay rate (one parameter). We computed the correlation between participants’ AQ score and each parameter, correcting for multiple comparisons separately for each parameter group. Only a negative correlation between AQ and the zero-cost second-thought probability was marginally significant (*r_s_* = −.22, *p* = .07, uncorrected *p* = .023), which suggests that higher AQ participants were less likely to use the second stage to reconsider stopping sampling in the zero-cost conditions, where the optimal strategy was to sample as many as possible. Though intuitive and consistent with the AQ effects on efficiency, we found this correlation would vanish when only the participants who were best fit by the 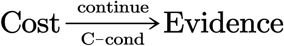 model were included (*r_s_* = −.02, *p* = .86) and thus might have been an epi-phenomenon arising from different individuals’ different decision strategies.

Next we tested whether participants’ autistic traits influenced the decision strategies they used. As shown in our results of model comparisons, participants may have used a variety of different two-stage decision processes: Among the 104 participants, 52 participants were best fit by the 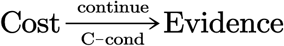 model and the remaining participants by the other two-stage models. It is also possible that the same individual may have used different decision processes in different choices. The assumptions of the 12 two-stage models, as we specified earlier, differed in three dimensions. On each dimension, we could classify the 12 models into different families (e.g. cost-first vs. evidence-first models concerning which factor controls the first stage). We quantified a specific participant’s decision strategies on the dimension by the participant’s mean AICc difference between the different families of models and computed its correlation with AQ (corrected for possible multiple comparisons on the dimension). We found that the AICc difference between cost-first and evidence-first model families (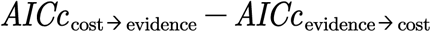) was negatively correlated with AQ (*r_s_* = −.23, *p* = .018; Fig 6a). An alternative analysis using the tripartite division of participants into AQ groups showed similar results (S1 Fig). Little correlations were found between 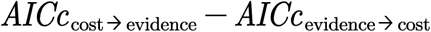 and other demographic variables including IQ, age, and gender (S6 Fig).

**Fig 6.**
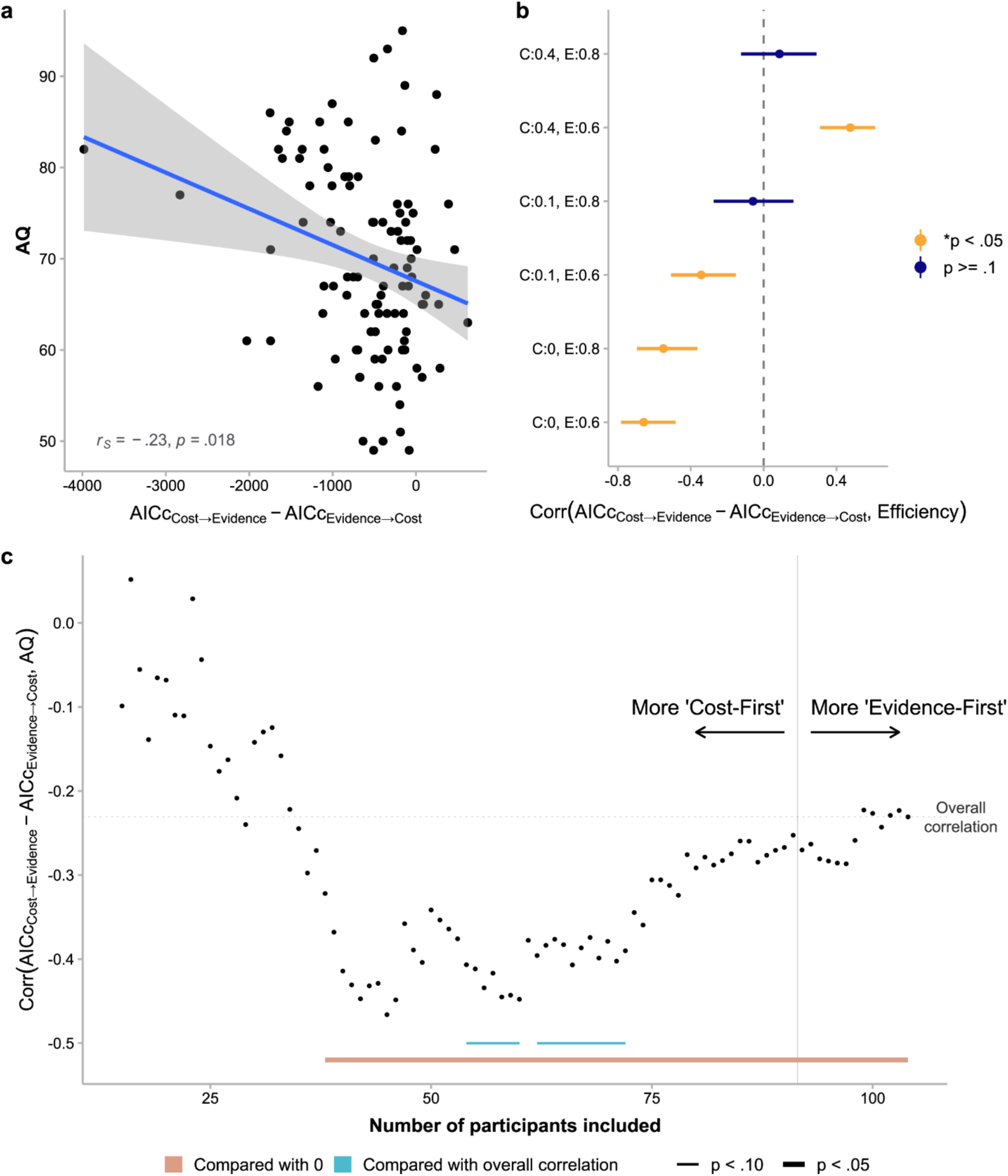
Effects of autistic traits on decision process and how it relates to sampling optimality. (a) Correlation between AQ and 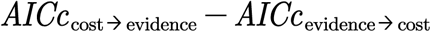. More positive 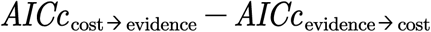 indicates stronger preference for cost-first over evidence-first decision processes, while more negative 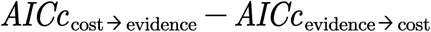 indicates the reverse. Each dot is for one participant. The blue line and the shaded area respectively represent regression line and standard error. (b) Correlation coefficients between 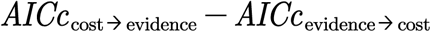 and efficiency for each cost and evidence condition. C:0 = zero-cost, C:0.1 = low-cost, C:0.4 = high-cost, E:0.6 = low-evidence, E:0.8 = high-evidence. Error bars represent FDR-corrected 95% confidence intervals. All these correlations were consistent with what we would expect if AQ influences sampling efficiency through its influence on the use of cost-first vs. evidence-first decision processes. For example, given that AQ was negatively correlated with 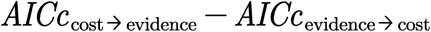, and 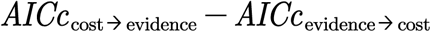 was negatively correlated with the efficiency in the zero-cost, low-evidence condition, we would expect AQ to be positively correlated with the efficiency in the zero-cost, low-evidence condition, and indeed it was. (c) Correlation between AQ and 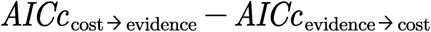 varied with the value of 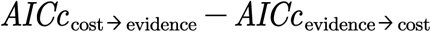 . We ranked all participants by 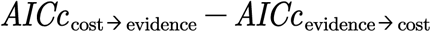 in ascending order, that is, from the strongest preference for cost-first to the strongest preference for evidence-first, and plot the Spearman’s correlation coefficient between 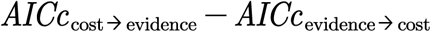 and AQ as a function of the number of participants included in the correlation analysis. The observed overall negative correlation and the stronger correlation given only the cost-first-dominated participants were included supports the cost-first vs. balanced-strategy hypothesis (see text): Participants with higher AQ tended to always consider cost first, while those with lower autistic traits considered cost or evidence first in a more balanced way. Statistical significance marked on the plot was based on cluster-based permutation tests (see Methods).

We assured that such differences in decision process could cause the observed autistic trait-related effects in sampling optimality by computing the correlation between 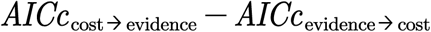 and efficiency for each cost and evidence condition (corrected for 6 comparisons). The correlation (Fig 6b) was significantly negative for the zero-cost, low-evidence condition (*r_s_* = −.66, *p* < .001), the zero-cost, high-evidence condition (*r_s_* = −.55, *p* < .001), and the low-cost, low-evidence condition (*r_s_* = −.34, *p* < .001), and was significantly positive for the high-cost, low-evidence condition (*r_s_* = .48, *p* < .001). All these correlations were consistent with what we would expect if AQ influences sampling efficiency through its influence on the use of cost-first vs. evidence-first decision processes. For example, given that AQ was negatively correlated with 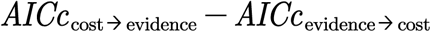, and 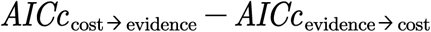 was negatively correlated with the efficiency in the zero-cost, low-evidence condition, we would expect AQ to be positively correlated with the efficiency in the zero-cost, low-evidence condition, and indeed it was. Similar correlations were also found between 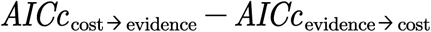 and sampling bias 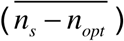 or sampling variation (*SD*(*n_s_*)) (S7 Fig).

Given that all participants were either much better modeled by cost-first models (i.e. 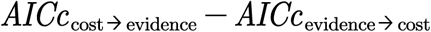 ≪ 0) or almost equivalently well by cost-first and evidence-first models (i.e. 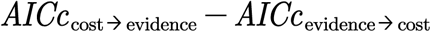 ≈ 0) (Fig 6a), the negative correlation between 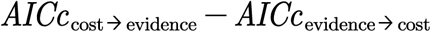 and AQ implies that participants with higher AQ preferred to consider cost first, while those with lower AQ preferred to have cost-first and evidence-first decisions more balanced (instead of preferring evidence first). If this cost-first vs. balanced-strategy (instead of cost-first vs. evidence-first) hypothesis for higher vs. lower AQ is true, we would also expect the correlation between 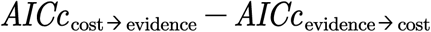 and AQ to be weak for those whose decisions were almost equally likely to be cost-first or evidence-first (i.e. 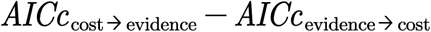 ≈ 0). In other words, we expect the correlation to be stronger if only the participants whose decisions were more dominated by cost-first (i.e. 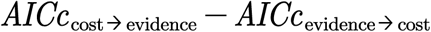 ≪ 0) is included. To test this, we ranked all participants by 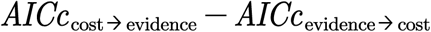 in ascending order and plot the Spearman’s correlation coefficient between 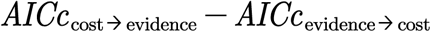 and AQ as a function of the number of participants included in the correlation analysis (Fig 6c). The correlation was statistically significant when the number of participants included was large enough (cluster-based permutation test, *p* = .006). In addition, compared to the overall correlation across the 104 participants, the correlation indeed appeared stronger when only the cost-first-dominated participants were included, which reached marginal significance when the number of participants included was between 54 and 60 (cluster-based permutation test, *p* = .083) or between 65 and 72 (*p* = .081). This provides further evidence for the cost-first vs. balanced-strategy hypothesis and suggests that participants with different levels of autistic traits differ in the diversity of their decision processes: Participants with higher AQ tended to always consider cost first, while those with lower autistic traits considered cost or evidence first in a more balanced way.

In the two-stage decision process we modeled, because the second stage is only probabilistically recruited, factors considered in the first stage would effectively leverage a greater influence on the sampling choice than those of the second stage. In other words, always being cost-first means the sampling choice is mainly determined by cost-related factors, while sometimes cost-first and sometimes evidence-first means the sampling choice is more of a tradeoff between cost- and evidence-related factors. Neither strategy is necessarily optimal but may approximate the optimal strategy in different situations: The former is closer to optimal when the optimal strategy does not depend on evidence, while the latter is closer to optimal when the optimal strategy varies with both cost and evidence. Participants’ differences in strategic diversity thus explain the autistic trait-related differences we observed in efficiency.

## Discussion

Humans must sample the environment properly to balance the advantage of gaining additional information against the cost of time, energy, and money [50]. Previous research suggests that suboptimal information sampling may be a fundamental deficit in ASD [4,14–17,51]. In the current study, we tested healthy adults with different levels of autistic traits to investigate how autistic traits influence information sampling decisions. We found that participants adjusted their sample sizes according to both sampling cost and evidence gain and were overall close to optimality. However, there were also systematic deviations from optimality which varied with levels of autistic traits. Computational modeling allowed us to characterize the decision process of sampling choices by two stages. The two-stage model well predicted the bimodality of DT distributions as well as the positive correlation between mean DT and the second-thought probability estimated from sampling choices. Autistic traits influenced the strategic diversity concerning whether cost or evidence is considered first.

Previous ASD studies that had used similar bead-sampling tasks yielded inconclusive results: One study found that adolescents with ASD sampled more than the control group [52], whereas a second study of adults with ASD found the reverse [40]. As to healthy people with higher autistic traits, we did not find overall oversampling or undersampling but more subtle differences. To ask whether people with ASD or higher autistic traits oversample or undersample information is probably not a proper question. In fact, both oversampling and undersampling may lower one’s expected gain, depending on the rewarding structure of the environment. As we suggested in the Introduction, a more important question is whether autistic traits influence one’s ability to sample optimally, that is, to balance sampling cost and information gain. In previous ASD studies [40,52], sampling incurred no explicit cost but implicit cost such as time or cognitive effort whose exact value to a specific individual is hard to measure, therefore we could hardly compare the optimality of different individuals’ performances. By introducing explicit monetary cost for sampling (as Juni et al. did [50]) in our experiment, we were able to evaluate sampling cost as a potential moderator for autistic trait-related differences in information sampling. Indeed, we found that people with higher autistic traits can be more optimal or less optimal than those with lower autistic traits depending on the level of sampling cost.

Sevgi, Diaconescu, Tittgemeyer, and Schilbach [53] demonstrated how computational modeling can be a powerful tool in deepening our understanding of autistic-trait-related cognitive processes and separating the affected processes from the intact ones. They found that autistic traits do not, as usually believed, influence individuals’ ability of learning social cues but only influence the weight assigned to social cues in decision making.

Similarly, the autistic-trait-related differences in sampling decisions we found through computational modeling are surprisingly selective. Participants with different levels of autistic traits were indistinguishable in their ability to weigh sampling cost or evidence gain in the two decision stages. What distinguished them was the strategic diversity across choices concerning whether to consider cost or evidence in the first stage. Participants with higher autistic traits were less diverse and stuck more to evaluating cost first.

Studies using autistic traits as a surrogate for studying ASD have revealed congruent and converging autistic-trait-related effects as those of ASD [9,10,53–57]. Although our findings could provide some insights on how autistic traits could influence people’s information sampling, we should also be aware that high autistic traits in typical people are not equivalent to symptoms of ASD [58–60] and autistic-trait-related differences do not necessarily characterize the differences between people with and without ASD. Thus, future research should test people with ASD to see how their information sampling differs from the typical population.

In our task, information sampling is instrumental—additional information would increase the probability of correct judgment. There are also situations where information is non-instrumental, for example, the information that is gathered after one’s decision and that would not change the outcome of the decision. Both humans [30–35] and non-human primates [36–39] are willing to pay for non-instrumental information, especially when it is good news. Whether autistic traits influence one’s tendency to seek non-instrumental information is a question for future research.

To summarize, we find that people with different levels of autistic traits differ in the optimality of information sampling and these differences are associated with their strategic diversity in the decision process. Recent studies suggest that autistic traits may influence an individual’s ability of adaptively using her own information processing capability while not influencing the capability itself. For example, autistic traits may only influence the tendency to use social information but not the capability to perceive it [53], or may only influence the flexibility of updating learning rate but not probabilistic learning itself [10]. Our results add to this line of findings that autistic-trait-related differences may come from differences in higher-level cognitive functions other than primary information processing.

## Methods

### Ethics Statement

The experiment had been approved by the Institutional Review Board of School of Psychological and Cognitive Sciences at Peking University (#2016-03-03). All participants provided written informed consent and were paid for their time plus performance-based bonus.

### Experiment

#### Participants

One hundred and fourteen college student volunteers participated in our experiment. Ten participants were excluded. Six of them were IQ outliers, one misunderstood instructions, one had a strong judgment bias towards one type of stimuli, one did not draw any bead in 286/288 of the trials, and one had a poor judgment consistency. This resulted in a final sample size of 104 participants (42 males, aged 18-28).

We estimated effect size a priori based on a mini meta-analysis of previous literature [61] on autistic-trait-related perceptual or cognitive differences [9,53–55,57,62–65], which was *r_s_* = .36. To achieve a statistical power of 0.80 under the significance level of .05, we would require 57 participants. However, considering initial effect sizes are often inflated [66], we doubled the estimate and sought to test around 114 participants with some attrition expected.

#### IQ test

Combined Raven Test (CRT) was used to measure participants’ IQ for control purpose. Raw CRT scores of all 114 participants averaged 67.69 (s.d., 4.71) and ranged from 41 to 72. Six of the participants (scoring from 41 to 58) fell out of two standard deviations of the mean and was excluded from further analyses along with four other participants (as mentioned above). The remaining 104 participants had a mean CRT score of 68.65 (s.d., 2.82; ranging from 61 to 72), corresponding to a mean IQ score of 117.68.

#### AQ test

Autism Spectrum Quotient (AQ) questionnaire [18] was used to quantify participants’ autistic traits. AQ questionnaire is a 4-point self-reported scale with 50 items measuring five type of autistic characteristics: social interaction, attentional switch, attention to detail, imagination, and communication. Though the 4-point scale was sometimes reduced to binary coding [18], we adopted the full 4-point scoring system (“definitely disagree”, “slightly disagree”, “slightly agree”, “definitely agree” respectively scored 0–3) to maximize the coverage of latent autistic traits [25,67–69].

The AQ scores of the 104 participants were normally distributed (Shapiro-Wilk normality test, *W* = 0.99, *p* = .32; S8 Fig) with mean 69.97 and standard deviation 10.48, ranging from 49 to 95. There was little correlation between AQ and IQ, *r_s_* = −.01, *p* = .95, AQ and age, *r_s_* = −.08, *p* = .40, or AQ and gender, biserial correlation *r_s_* = .13, *p* = .31.

#### Apparatus

All stimuli of the bead-sampling task were visually presented on a 21.5-inch computer screen controlled by MATLAB R2016b and PsychToolbox [70–72].

Participants were seated approximately 60 cm to the screen. Responses were recorded via the keyboard.

#### Procedure

On each trial of the experiment (Fig 1a), participants saw a pair of jars on the left and right of the screen, each containing 200 pink and blue beads. The pink-to-blue ratios of the two jars were either 60%:40% vs. 40%:60%, or 80%:20% vs. 20%:80%. Participants were told that one jar had been secretly selected, and their task was to infer which jar was selected. Each time they pressed the space bar, one bead was randomly sampled with replacement from the jar and presented on the screen, appended to the end of the sampled bead sequence. Participants were free to draw 0 to 20 bead samples, but each sample might incur a cost. The cost per sample on each trial could be 0, 0.1, or 0.4 points. A green bar on the top of the screen indicated how many bonus points remained (10 points minus the total sampling cost by then). When participants were ready for inference, they pressed the Enter key to quit sampling and judged whether the pre-selected jar was the left or right jar by pressing the corresponding arrow key. Feedback followed immediately. If their judgment was correct, participants would receive the remaining bonus points; otherwise nothing. Bonus points accumulated across trials and would be converted into monetary bonus after the experiment. Participants were encouraged to sample wisely to maximize their winning.

The pink-dominant jar was pre-selected on half of the trials and the blue-dominant jar on the other half. Their left/right positions were also counterbalanced across trials. In the formal experiment, the two evidence (i.e. bead ratio) conditions (60/40 and 80/20) were randomly mixed within each block and the three cost conditions (0, 0.1, and 0.4) were blocked. Besides being visualized by the green bar on each trial, cost for each block was also informed at the beginning of the block. The order of cost blocks was counterbalanced across participants. We further confirmed that block order (6 permutations) had no significant effects on participants’ sampling choices (efficiency: *F*_5,97.90_ = 2.06, *p* = .08, 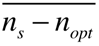 : *F*_5,97.99_ = 1.51, *p* = .19, *SD*(*n_s_*) : *F*_5,97.97_ = 1.53,*p* = .19) or decision times (*F*_5,98_ = 0.60 *p* = .70). Each of the six conditions was repeated for 48 times, resulting in 288 trials. The formal experiment was preceded by 24 practice trials. Participants first performed the experiment, then the Combined Raven Test and last the AQ questionnaire, which took approximately 1.5 hours in total.

### Statistical Analyses

All statistical analyses (except for group-level Bayesian model comparison) were conducted in R 3.5.3 [73].

#### Linear mixed models (LMMs)

Linear mixed models were estimated using “afex” package [74], whose *F* statistics, degrees of freedom of residuals (denominators), and *p*-values were approximated by Kenward-Roger method [75,76]. Specifications of random effects followed parsimonious modeling [77]. For significant fixed effects, “emmeans” package was used to test post hoc contrasts [78]. Interaction contrasts were performed for significant interactions and, when higher order interactions were not significant, pairwise or consecutive contrasts were performed for significant main effects. Statistical multiplicity of the contrasts was controlled by a single-step adjustment, which used multivariate *t* distributions to estimate the critical value for conducted contrasts [79,80].

LMM1: decision efficiency is the dependent variable; fixed effects include an intercept, the main and interaction effects of AQ, cost, and ratio (evidence); random effects include correlated random slopes of costs and ratios within participants and random participant intercept.

LMM2: sampling bias (mean number of actual sampling minus optimal number of sampling 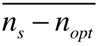) is the dependent variable; the fixed and random effects are the same as LMM1.

LMM3: standard deviation of the number of sampling (*SD*(*n_s_*)) is the dependent variable; the fixed and random effects are the same as LMM1.

LMM4: mean decision time (DT) across all sampling choices of a condition is the dependent variable; the fixed and random effects are the same as LMM1.

LMM5: DT of each sample number (1 to 20 samples) averaged over all trials is the dependent variable; fixed effects involve an intercept, the main and interaction effects of AQ and sample number, and random effects include a random participant intercept. The model also incorporated weights on the residual variance for each aggregated data point to account for the different number of raw DTs for each sample number of each participant.

LMM6: the dependent variable is the same as LMM4; in addition to the fixed and random effects of LMM1, the linear effect of second-thought probability is included in the fixed effects, and a random slope of the second-thought probability that is uncorrelated with the random intercept is included in the random effects.

Following Jones et al. [81], we identified three “likely noncompliant” outlier observations in the number of bead samples for each condition based on nonparametric boxplot statistics, that is, those whose values were lower than the 1st quartile or higher than the 3rd quartile of all the observations in the condition by more than 1.5 times of the interquartile range (see S9 Fig). These noncompliant observations (not participants per se) were excluded from LMMs 1–3.

To examine possible non-linear effects of AQ, we constructed LMMs that included AQ^2^ and its interaction with cost and ratio as additional fixed-effects terms separately for LMM1–6. We found that adding the second order terms of AQ did not significantly improve the goodness-of-fit of any LMM.

#### Decision times (DTs)

Because stopping sampling involved a different key press, only DTs for continuing sampling were analyzed. Before any analysis of DTs, outliers of log-transformed DTs were excluded based on nonparametric boxplot statistics, with data points lower than the 1st quartile or higher than the 3rd quartile of all the log-transformed DTs by more than 1.5 times of the interquartile range defined as outliers.

#### Correlation analyses based on modeling results

Spearman’s rank correlations (denoted *r*) were computed between AQ and model measures (model parameter or model evidence), and between model measures and behavioral measures (efficiency, 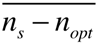, or *SD*(*n_s_*)). Except for the statistics in Fig 6c, multiple correlation tests were corrected using false discovery rate (FDR) to avoid the inflation of false alarm rates with multiple comparisons.

To test whether the curve of correlation coefficients between 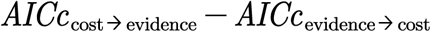 and AQ in Fig 6c was significantly different from 0 or the overall correlation at some points, we performed cluster-based permutation tests [82] as follows. For the test against 0, we first identified points that were significantly different from 0 at the uncorrected significance level of .05 using *t* tests and then grouped adjacent same-signed significant correlations into clusters. For each cluster, the absolute value of the summed Fisher’s z values transformed from *r*was defined as the cluster size. We randomly shuffled the values of 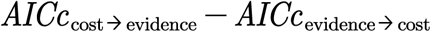 across participants to generate virtual data, calculated the correlation curve and recorded the maximum size of its clusters for the virtual data. This procedure was repeated for 10,000 times to produce a distribution of chance-level maximum cluster sizes, based on which we calculated the *p* value for each cluster in real data.

For the test against the overall correlation of 104 participants, we randomly shuffled the order of inclusion across participants and identified points that were significantly different from the overall correlation at the uncorrected significance level of .05 using Monte Carlo methods. Otherwise the permutation test was identical to that described above.

### Modeling

#### Expected gain

Given a specific sequence of bead samples, an ideal observer would always judge the preselected jar to be the one whose dominant color is the same as that of the sample sequence. In the case of a tie, the observer would choose the two jars with equal probability. Suppose the sample size is *n*, the maximal reward is 10 points, the unit sampling cost is *c*, and the percentage of predominated beads in the preselected jar is *q*. The expected probability of correct judgment is:

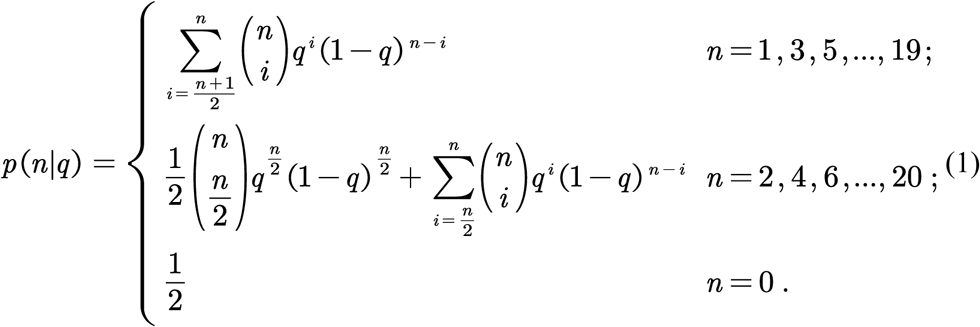

The expected gain is E[*Gain*|*n, q, c*] = (10 – *nc*)*p*(*n*|*q*). For a specific cost and evidence condition, the optimal sample size is the value of *n* that maximizes E[*Gain*|*n, q, c*].

#### One-stage models

We modeled participants’ each choice of whether to continue or stop sampling (i.e. whether to press the space bar or Enter key) as a Bernoulli random variable, with the probability of stopping sampling determined by cost- or evidence-related factors. Pressing the Enter key after 20 samples was not included as a choice of stopping sampling, because participants had no choice but to stop by then.

We considered two families of models: one-stage and two-stage models. The description for each model is summarized in S1 Table. In one-stage models, the probability of stopping sampling on the *i*-th trial after having drawn *j* beads is determined by a linear combination of *K* decision variables (DVs) via a logistic function:

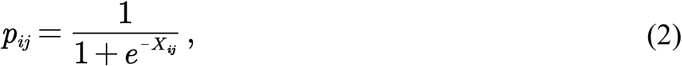

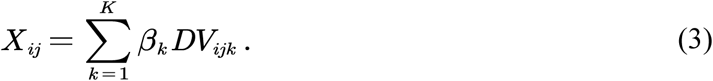

Different one-stage models differed in whether cost-related variables, evidence-related variables, or both served as DVs (S1 Table).

Cost-only one-stage model (denoted *Cost only*): cost-related variables as DVs, including unit cost per bead (categorical: 0, 0.1, or 0.4), number of beads sampled (*j*), and total sampling cost (product of the former two DVs).

Evidence-only without decay one-stage model (denoted *Evidence only w/o decay*): evidence-related variables as DVs, including unit log evidence per bead (i.e., ln(60/40) or ln(80/20)), absolute value of cumulative information (cumulative information refers to the difference between the numbers of pink and blue bead samples), total log evidence (product of the former two DVs), and the correctness and the number of bead samples in last trial.

Cost + evidence without decay one-stage model (denoted *Cost + Evidence w/o decay*): both cost-related and evidence-related variables as DVs.

Cost + evidence with decay one-stage model (denoted *Cost + Evidence*): both cost-related and decayed evidence-related variables as DVs.

In models with decayed evidence, cumulative information (CI) is modulated by a decay parameter α:

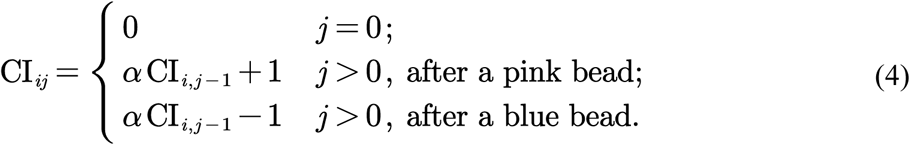

The DVs of absolute value of cumulative information and total log evidence in the models with decay are modulated by the decay parameter accordingly.

#### Two-stage models

In two-stage models, sampling choices may involve two decision stages, with the probability of reaching the decision of stopping sampling in each stage being

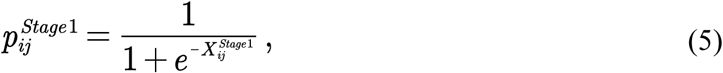

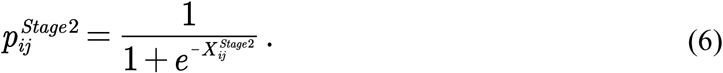

Whether to enter the second stage is probabilistic, conditional on the decision reached in the first stage. For models where the second stage is triggered by the decision of continuing sampling in the first stage, the overall probability of stopping sampling can be written as:

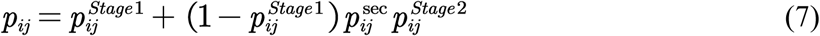

Here denotes second-thought probability—the probability of using the second stage given that the first stage concludes with continuing sampling, whose value is defined differently in different models as specified below. Alternatively, for models where the second stage is triggered by the decision of stopping sampling in the first stage, the overall probability of stopping sampling can be written as:

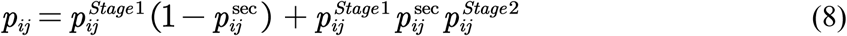

Each stage works in the same way as one-stage models do (Eqs. 2–4) and is influenced by mutually exclusive sets of DVs (S1 Table). We considered two-stage models whose assumptions differ in three dimensions: (1) which factors control the first stage and which control the second stage (cost-first or evidence-first), (2) what kind of decision in the first stage (continuing or stopping sampling) has a chance to trigger the second stage, and (3) what determines the probability to enter the second stage (“second-thought probability”) after a qualified first-stage decision (the cost condition, the evidence condition, or the probability of stopping in the first-stage decision). A full 2×2×3 combinations resulted in 12 different two-stage models. The assumptions for each dimension are specified below.

Cost-first two-stage models (models denoted by 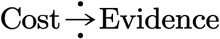): cost-related variables as first-stage DVs and decayed evidence-related variables as second-stage DVs.

Evidence-first two-stage models (models denoted by 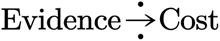): decayed evidence-related variables as first-stage DVs and cost-related variables as second-stage DVs.

Continue-then-2nd-thought two-stage models (models denoted by 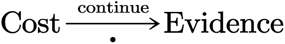 or 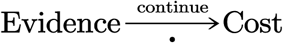): If stopping sampling is the decision in the first stage, it is finalized and there is no second stage; otherwise, either continuing sampling becomes the final decision, or the decision is re-evaluated in the second stage.

Stop-then-2nd-thought two-stage models (models denoted by 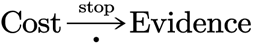 or 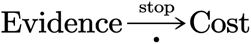): If continuing sampling is the decision in the first stage, it is finalized and there is no second stage; otherwise, either stopping sampling becomes the final decision, or the decision is re-evaluated in the second stage.

Cost-controls-2nd-thought two-stage models (models denoted by 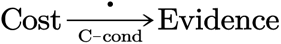 or 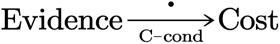): The second-thought probability is controlled by the cost condition, with 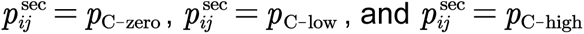, respectively for the zero-, low-, and high-cost conditions, where *p*_C-zero_, *p*_C-low_ and *p*_C-high_ are free parameters.

Evidence-controls-2nd-thought two-stage models (models denoted by 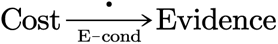 or 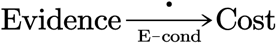): The second-thought probability is controlled by the evidence condition, with 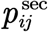 = *p*_E-low_ and 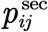 = *p*_E-high_ respectively for the low- and high-evidence conditions, where *p*_E-low_ and *p*_E-high_ are free parameters.

Flexible-2nd-thought two-stage models (models denoted by 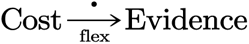 or 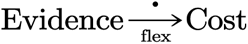): The second-thought probability is a function of the probability of stopping sampling in the first stage,

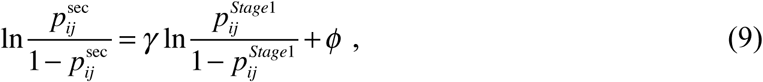

where *γ* and *ϕ*are free parameters.

The intuition behind this form of second-thought probability is that participants should be likely to use the second stage to stop sampling when they are reluctant to continue but end up with choosing continue in the first stage, and likewise for the reverse case.

For both one- and two-stage models, given that the probability of stopping sampling on the *i*-th trial after having drawn *j* beads is *p_ij_*, the likelihood of observing a specific choice *c_ij_* (0 for continue and 1 for stop) is

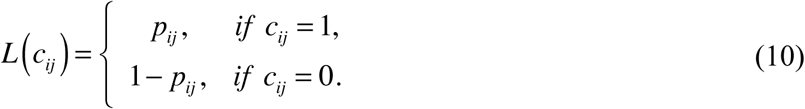

#### Modeling decision times (DTs)

Evidence-accumulation models are the common practice to model the response time (RT) of human decision-making, which can capture the three properties of the observed RT distributions [83]: (1) RT distributions are positively skewed; (2) More difficult choices (i.e. when the two options are more closely matched in the probability of being chosen) lead to longer RTs. (3) Correct choices (i.e. choosing the option with the higher value) can have equal, shorter, or longer RTs than wrong choices (i.e. choosing the option with the lower value). However, evidence-accumulation models would be computationally intractable if applied to the two-stage decision process of our interest, because there have been no analytical form or efficient numerical algorithms to deal with the RT distribution resulting from two evidence-accumulation processes, especially when the variables controlling each evidence-accumulation process vary from choice to choice, as in our case.

Therefore, we modeled participants’ decision time (DT) for each sampling with a simplified form that is able to capture the three properties summarized above. For one-stage models or the first stage of two-stage models, we have

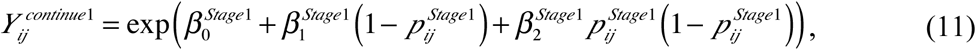

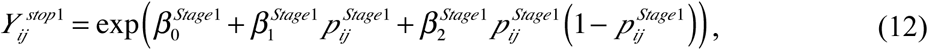

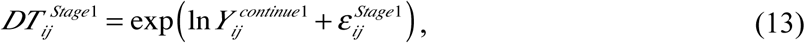

where 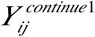 and 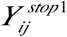 denote the expected DTs respectively for continuing and stopping sampling, which have the same form expect that the 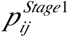 in Eq. 11 is replaced by 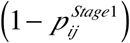 in Eq. 11. 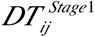 denotes the observed DT if the decision of continuing sampling is made in the first stage. Here 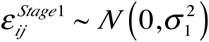 is a Gaussian noise term so that 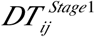 is log-normally distributed, satisfying Property (1). The quadratic term, 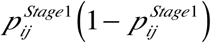, allows 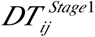 to vary with choice difficulty so as to satisfy Property (2). The inclusion of the 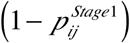 term, would enable the three possibilities of Property (3). The 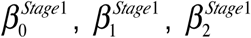, and 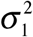 are free parameters.

The expected total DT of reaching the decision of continuing sampling in the second stage equals to the time required by the first stage plus that of the second stage and has the forms

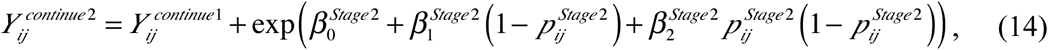

and

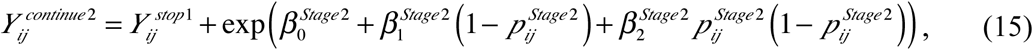

respectively for continue-then-2nd-thought and stop-then-2nd-thought models. The observed DT of continuing sampling in the second stage is then

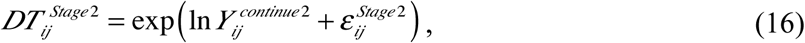

where 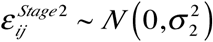 is a Gaussian noise term. The 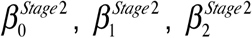, and 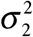 are free parameters.

Thus, for one-stage models, the likelihood of observing a specific *DT_ij_* for drawing the *(j+1)*-th bead on the *i*-th trial is

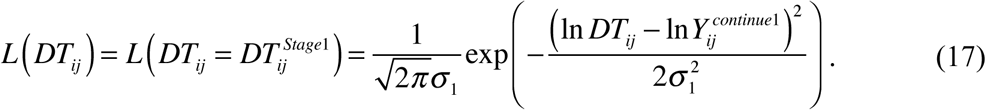

For two-stage models, where *DT_ij_* is a mixture of 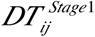 and 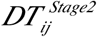, its likelihood follows

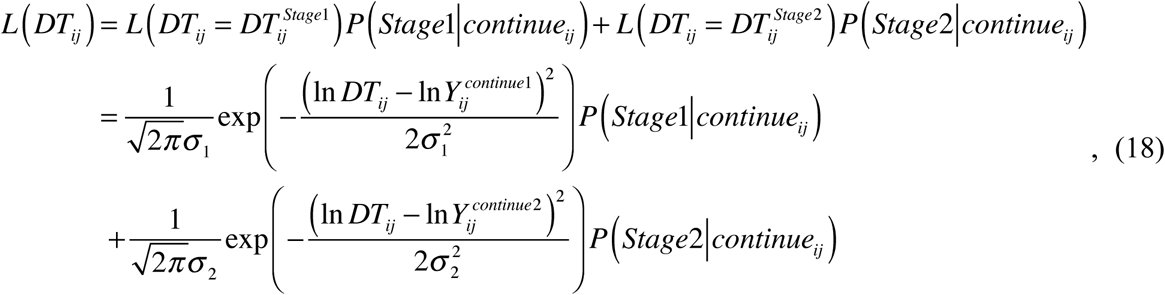

where *P* (*Stage*1 | *continue_ij_*) and *P* (*Stage2 | continue_ij_*) respectively refer to the probabilities that the choice is finalized at Stage 1 and Stage 2, given that continuing sampling is the choice. These probabilities are computed based on the corresponding choice model, which are

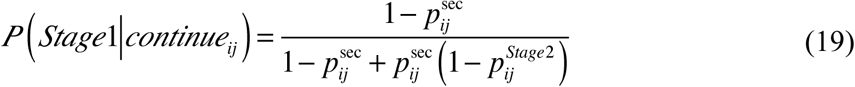

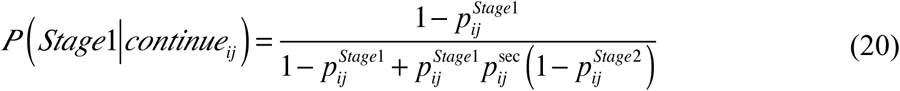

and respectively for continue-then-2nd-thought and stop-then-2nd-thought two-stage models, and

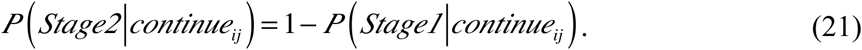

The 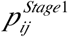, 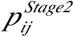, are 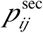 defined earlier in the choice model and estimated from participants’ choices.

#### Joint log likelihood of choice and DT

For a specific sampling choice modeled by two-stage models, the likelihood of the joint observation of *continue_ij_* and *DT_ij_* is

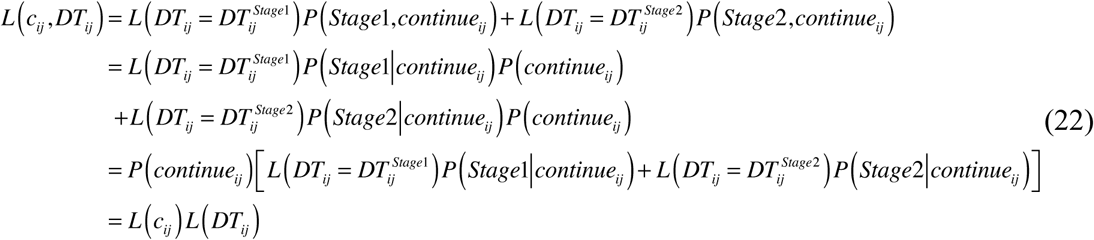

That is, the joint likelihood is equivalent to the product of the likelihoods of choice (Eq. 10) and DT (Eqs. 17-18). The same equivalence holds for one-stage models, whose proof is a special case of that of two-stage models. For the joint log likelihood summed over trials, we have

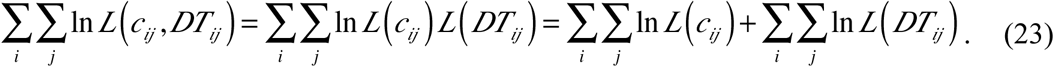

Therefore, we used the sum of the log likelihoods of the choice and DT models for model comparisons.

#### Model fitting

Each one- or two-stage model consists of two parts: choice and DT. We first fit each choice model separately for each participant to the participant’s actual sampling choices using maximum likelihood estimates. As an example, if the participant samples 5 beads on a trial, she has a sequence of 6 binary choices on the trial (000001, with 0 for continue and 1 for stop). Different models differ in how the likelihood of generating a specific choice (0 or 1) varies with the cost or evidence observed before the choice. For one-stage models, where all decision variables control the choice in one stage, the influence of cost- or evidence-related variables is fixed across experimental conditions. In contrast, for two-stage models, the decision variables that control the second stage exert variable influences on the choice, because the probability for the second stage to be recruited varies with experimental conditions. The observed choice patterns in the experiment thus allowed us to discriminate different models, including one- and two-stage models.

For a specific fitted choice model, we could compute the second-thought probability, whenever applicable, as well as the probabilities of choosing stopping at each stage. With this information, we then fit the corresponding DT model to the participant’s DTs to estimate the DT-unique parameters.

We chose to optimize the parameters of choice and DT models in this way instead of optimizing them simultaneously to avoid the computational intractability of fitting a large number of parameters. In addition, choices and DTs can serve as independent tests for the two-stage decision process we proposed.

All coefficients *β_k_* of decision variables, second-thought probabilities 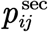, decay parameter *α*, and all *β* and *σ* in DT models were estimated as free parameters using maximum likelihood estimates. All parameters were unbounded, except that 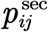 of cost-controlled and evidence-controlled second-thought models and *α* were bounded to [0, 1], and 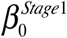, 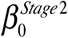 *σ*_1_, and *σ*_2_ of DT models were bounded to (0, Inf). Optimization was implemented by the *fmincon* function with interior-point algorithm in MATLAB R2017a.

#### Model comparison

The Akaike Information Criterion corrected for small samples (AICc) [44,45] and Bayesian Information Criterion (BIC) were calculated as model evidence for model comparison. In the computation of these information measures, the number of “trials” of a participant’s dataset was defined as the number of DTs modeled for the participant. The ΔAICc (ΔBIC) for a specific model was computed for each participant as the AICc (BIC) difference between the model and the participant’s best-fitting model (i.e. the model with the lowest AICc (BIC)). The summed ΔAICc (ΔBIC) across participants was used for fixed-effects comparisons. Group-level Bayesian model selection [46,47] was used to provide an omnibus measure across individual participants that takes into account random effects.

## References

1. Dall S, Giraldeau L, Olsson O, Mcnamara J, Stephens D. Information and its use by animals in evolutionary ecology. Trends Ecol Evol. 2005;20: 187–193. doi:10.1016/j.tree.2005.01.010

2. Stephens DW, Krebs JR. Foraging Theory. Princeton University Press; 1986.

3. American Psychiatric Association. Diagnostic and Statistical Manual of Mental Disorders (DSM-5®). American Psychiatric Pub; 2013.

4. Palmer CJ, Lawson RP, Hohwy J. Bayesian approaches to autism: Towards volatility, action, and behavior. Psychol Bull. 2017;143: 521–542. doi:10.1037/bul0000097

5. Au□Yeung SK, Kaakinen JK, Benson V. Cognitive Perspective-Taking During Scene Perception in Autism Spectrum Disorder: Evidence From Eye Movements. Autism Res. 2014;7: 84–93. doi:10.1002/aur.1352

6. Song Y, Hakoda Y, Sang B. A selective impairment in extracting fearful information from another’s eyes in Autism. Autism Res. 2016;9: 1002–1011. doi:10.1002/aur.1583

7. Chambon V, Farrer C, Pacherie E, Jacquet PO, Leboyer M, Zalla T. Reduced sensitivity to social priors during action prediction in adults with autism spectrum disorders. Cognition. 2017;160: 17–26. doi:10.1016/j.cognition.2016.12.005

8. Goris J, Braem S, Nijhof AD, Rigoni D, Deschrijver E, Cruys SV de, et al. Sensory Prediction Errors Are Less Modulated by Global Context in Autism Spectrum Disorder. Biol Psychiatry Cogn Neurosci Neuroimaging. 2018;0. doi:10.1016/j.bpsc.2018.02.003

9. Lawson RP, Aylward J, Roiser JP, Rees G. Adaptation of social and non-social cues to direction in adults with autism spectrum disorder and neurotypical adults with autistic traits. Dev Cogn Neurosci. 2018;29: 108–116. doi:10.1016/j.dcn.2017.05.001

10. Lawson RP, Mathys C, Rees G. Adults with autism overestimate the volatility of the sensory environment. Nat Neurosci. 2017;20: nn.4615. doi:10.1038/nn.4615

11. Manning C, Tibber MS, Charman T, Dakin SC, Pellicano E. Enhanced Integration of Motion Information in Children With Autism. J Neurosci. 2015;35: 6979–6986. doi:10.1523/JNEUROSCI.4645-14.2015

12. Palmer CJ, Paton B, Kirkovski M, Enticott PG, Hohwy J. Context sensitivity in action decreases along the autism spectrum: a predictive processing perspective. Proc R Soc Lond B Biol Sci. 2015;282: 20141557. doi:10.1098/rspb.2014.1557

13. Turi M, Burr DC, Igliozzi R, Aagten-Murphy D, Muratori F, Pellicano E. Children with autism spectrum disorder show reduced adaptation to number. Proc Natl Acad Sci U S A. 2015;112: 7868–7872. doi:10.1073/pnas.1504099112

14. Lawson RP, Rees G, Friston KJ. An aberrant precision account of autism. Front Hum Neurosci. 2014;8. doi:10.3389/fnhum.2014.00302

15. Pellicano E, Burr D. When the world becomes ‘too real’: a Bayesian explanation of autistic perception. Trends Cogn Sci. 2012;16: 504–510. doi:10.1016/j.tics.2012.08.009

16. van Boxtel JJA, Lu H. A predictive coding perspective on autism spectrum disorders. Front Psychol. 2013;4. doi:10.3389/fpsyg.2013.00019

17. van de Cruys S, Evers K, van der Hallen R, van Eylen L, Boets B, de-Wit L, et al. Precise minds in uncertain worlds: Predictive coding in autism. Psychol Rev. 2014;121: 649–675. doi:10.1037/a0037665

18. Baron-Cohen S, Wheelwright S, Skinner R, Martin J, Clubley E. The autism-spectrum quotient (AQ): Evidence from asperger syndrome/high-functioning autism, males and females, scientists and mathematicians. J Autism Dev Disord. 2001;31: 5–17.

19. Hoekstra RA, Bartels M, Verweij CJH, Boomsma DI. Heritability of Autistic Traits in the General Population. Arch Pediatr Adolesc Med. 2007;161: 372–377. doi:10.1001/archpedi.161.4.372

20. Lundström S, Chang Z, Råstam M, Gillberg C, Larsson H, Anckarsäter H, et al. Autism Spectrum Disorders and Autisticlike Traits: Similar Etiology in the Extreme End and the Normal Variation. Arch Gen Psychiatry. 2012;69: 46–52. doi:10.1001/archgenpsychiatry.2011.144

21. Robinson EB, Koenen KC, McCormick MC, Munir K, Hallett V, Happé F, et al. Evidence That Autistic Traits Show the Same Etiology in the General Population and at the Quantitative Extremes (5%, 2.5%, and 1%). Arch Gen Psychiatry. 2011;68: 1113–1121. doi:10.1001/archgenpsychiatry.2011.119

22. Ronald A, Hoekstra RA. Autism spectrum disorders and autistic traits: A decade of new twin studies. Am J Med Genet B Neuropsychiatr Genet. 2011;156: 255–274. doi:10.1002/ajmg.b.31159

23. Sucksmith E, Roth I, Hoekstra RA. Autistic Traits Below the Clinical Threshold: Re-examining the Broader Autism Phenotype in the 21st Century. Neuropsychol Rev N Y. 2011;21: 360–89. doi:http://dx.doi.org/10.1007/s11065-011-9183-9

24. Wheelwright S, Auyeung B, Allison C, Baron-Cohen S. Defining the broader, medium and narrow autism phenotype among parents using the Autism Spectrum Quotient (AQ). Mol Autism. 2010;1: 10. doi:10.1186/2040-2392-1-10

25. Bralten J, Hulzen KJ van, Martens MB, Galesloot TE, Vasquez AA, Kiemeney LA, et al. Autism spectrum disorders and autistic traits share genetics and biology. Mol Psychiatry. 2017; doi:10.1038/mp.2017.98

26. Ruzich E, Allison C, Smith P, Watson P, Auyeung B, Ring H, et al. Measuring autistic traits in the general population: a systematic review of the Autism-Spectrum Quotient (AQ) in a nonclinical population sample of 6,900 typical adult males and females. Mol Autism. 2015;6: 2. doi:10.1186/2040-2392-6-2

27. Moutoussis M, Bentall RP, El-Deredy W, Dayan P. Bayesian modelling of Jumping-to-Conclusions bias in delusional patients. Cognit Neuropsychiatry. 2011;16: 422–447. doi:10.1080/13546805.2010.548678

28. Hauser TU, Moutoussis M, Dayan P, Dolan RJ. Increased decision thresholds trigger extended information gathering across the compulsivity spectrum. Transl Psychiatry. 2017;7: 1296. doi:10.1038/s41398-017-0040-3

29. Hauser TU, Moutoussis M, Iannaccone R, Brem S, Walitza S, Drechsler R, et al. Increased decision thresholds enhance information gathering performance in juvenile Obsessive-Compulsive Disorder (OCD). PLOS Comput Biol. 2017;13: e1005440. doi:10.1371/journal.pcbi.1005440

30. Hunt LT, Rutledge RB, Malalasekera WMN, Kennerley SW, Dolan RJ. Approach-Induced Biases in Human Information Sampling. PLOS Biol. 2016;14: e2000638. doi:10.1371/journal.pbio.2000638

31. Charpentier CJ, Bromberg-Martin ES, Sharot T. Valuation of knowledge and ignorance in mesolimbic reward circuitry. Proc Natl Acad Sci. 2018;115: E7255–E7264. doi:10.1073/pnas.1800547115

32. Kobayashi K, Ravaioli S, Baranès A, Woodford M, Gottlieb J. Diverse motives for human curiosity. Nat Hum Behav. 2019; 1. doi:10.1038/s41562-019-0589-3

33. Clark L, Robbins TW, Ersche KD, Sahakian BJ. Reflection Impulsivity in Current and Former Substance Users. Biol Psychiatry. 2006;60: 515–522. doi:10.1016/j.biopsych.2005.11.007

34. Bennett D, Bode S, Brydevall M, Warren H, Murawski C. Intrinsic Valuation of Information in Decision Making under Uncertainty. PLOS Comput Biol. 2016;12: e1005020. doi:10.1371/journal.pcbi.1005020

35. Iigaya K, Story GW, Kurth-Nelson Z, Dolan RJ, Dayan P. The modulation of savouring by prediction error and its effects on choice. Uchida N, editor. eLife. 2016;5: e13747. doi:10.7554/eLife.13747

36. Bromberg-Martin ES, Hikosaka O. Midbrain Dopamine Neurons Signal Preference for Advance Information about Upcoming Rewards. Neuron. 2009;63: 119–126. doi:10.1016/j.neuron.2009.06.009

37. Bromberg-Martin ES, Hikosaka O. Lateral habenula neurons signal errors in the prediction of reward information. Nat Neurosci. 2011;14: 1209–1216. doi:10.1038/nn.2902

38. Blanchard TC, Hayden BY, Bromberg-Martin ES. Orbitofrontal Cortex Uses Distinct Codes for Different Choice Attributes in Decisions Motivated by Curiosity. Neuron. 2015;85: 602–614. doi:10.1016/j.neuron.2014.12.050

39. Stoll FM, Fontanier V, Procyk E. Specific frontal neural dynamics contribute to decisions to check. Nat Commun. 2016;7: 11990. doi:10.1038/ncomms11990

40. Jänsch C, Hare DJ. An Investigation of the “Jumping to Conclusions” Data-Gathering Bias and Paranoid Thoughts in Asperger Syndrome. J Autism Dev Disord. 2014;44: 111–119. doi:10.1007/s10803-013-1855-2

41. Huq SF, Garety PA, Hemsley DR. Probabilistic Judgements in Deluded and Non-Deluded Subjects. Q J Exp Psychol Sect A. 1988;40: 801–812. doi:10.1080/14640748808402300

42. Ratcliff R, Smith PL, Brown SD, McKoon G. Diffusion Decision Model: Current Issues and History. Trends Cogn Sci. 2016;20: 260–281. doi:10.1016/j.tics.2016.01.007

43. Wolfe JM, Palmer EM, Horowitz TS. Reaction time distributions constrain models of visual search. Vision Res. 2010;50: 1304–1311. doi:10.1016/j.visres.2009.11.002

44. Cavanaugh JE. Unifying the derivations for the Akaike and corrected Akaike information criteria. Stat Probab Lett. 1997;33: 201–208. doi:10.1016/S0167-7152(96)00128-9

45. Hurvich CM, Tsai C-L. Regression and time series model selection in small samples. Biometrika. 1989;76: 297–307. doi:10.1093/biomet/76.2.297

46. Rigoux L, Stephan KE, Friston KJ, Daunizeau J. Bayesian model selection for group studies — Revisited. NeuroImage. 2014;84: 971–985. doi:10.1016/j.neuroimage.2013.08.065

47. Stephan KE, Penny WD, Daunizeau J, Moran RJ, Friston KJ. Bayesian model selection for group studies. NeuroImage. 2009;46: 1004–1017. doi:10.1016/j.neuroimage.2009.03.025

48. Schwarz G. Estimating the Dimension of a Model. Ann Stat. 1978;6: 461–464. doi:10.1214/aos/1176344136

49. Burnham KP, Anderson DR. Multimodel Inference: Understanding AIC and BIC in Model Selection. Sociol Methods Res. 2004;33: 261–304. doi:10.1177/0049124104268644

50. Juni MZ, Gureckis TM, Maloney LT. Information sampling behavior with explicit sampling costs. Decision. 2016;3: 147–168. doi:10.1037/dec0000045

51. Sinha P, Kjelgaard MM, Gandhi TK, Tsourides K, Cardinaux AL, Pantazis D, et al. Autism as a disorder of prediction. Proc Natl Acad Sci. 2014;111: 15220–15225. doi:10.1073/pnas.1416797111

52. Brosnan M, Chapman E, Ashwin C. Adolescents with Autism Spectrum Disorder Show a Circumspect Reasoning Bias Rather than ‘Jumping-to-Conclusions.’ J Autism Dev Disord. 2014;44: 513–520. doi:10.1007/s10803-013-1897-5

53. Sevgi M, Diaconescu AO, Tittgemeyer M, Schilbach L. Social Bayes: Using Bayesian Modeling to Study Autistic Trait–Related Differences in Social Cognition. Biol Psychiatry. 2016;80: 112–119. doi:10.1016/j.biopsych.2015.11.025

54. Robic S, Sonié S, Fonlupt P, Henaff M-A, Touil N, Coricelli G, et al. Decision-Making in a Changing World: A Study in Autism Spectrum Disorders. J Autism Dev Disord. 2015;45: 1603–1613. doi:10.1007/s10803-014-2311-7

55. Turi M, Burr DC, Binda P. Pupillometry reveals perceptual differences that are tightly linked to autistic traits in typical adults. eLife. 2018;7: e32399. doi:10.7554/eLife.32399

56. van Boxtel JJA, Dapretto M, Lu H. Intact recognition, but attenuated adaptation, for biological motion in youth with autism spectrum disorder. Autism Res. 2016;9: 1103–1113. doi:10.1002/aur.1595

57. van Boxtel JJA, Lu H. Impaired Global, and Compensatory Local, Biological Motion Processing in People with High Levels of Autistic Traits. Front Psychol. 2013;4. doi:10.3389/fpsyg.2013.00209

58. Baghdadli A, Russet F, Mottron L. Measurement properties of screening and diagnostic tools for autism spectrum adults of mean normal intelligence: A systematic review. Eur Psychiatry. 2017;44: 104–124. doi:10.1016/j.eurpsy.2017.04.009

59. Ashwood KL, Gillan N, Horder J, Hayward H, Woodhouse E, McEwen FS, et al. Predicting the diagnosis of autism in adults using the Autism-Spectrum Quotient (AQ) questionnaire. Psychol Med. 2016;46: 2595–2604. doi:10.1017/S0033291716001082

60. Sizoo BB, Horwitz E, Teunisse J, Kan C, Vissers C, Forceville E, et al. Predictive validity of self-report questionnaires in the assessment of autism spectrum disorders in adults. Autism. 2015;19: 842–849. doi:10.1177/1362361315589869

61. Goh JX, Hall JA, Rosenthal R. Mini Meta-Analysis of Your Own Studies: Some Arguments on Why and a Primer on How: Mini Meta-Analysis. Soc Personal Psychol Compass. 2016;10: 535–549. doi:10.1111/spc3.12267

62. Karvelis P, Seitz AR, Lawrie SM, Seriès P. Autistic traits, but not schizotypy, predict increased weighting of sensory information in Bayesian visual integration. eLife. 2018;7: e34115. doi:10.7554/eLife.34115

63. Chouinard PA, Unwin KL, Landry O, Sperandio I. Susceptibility to Optical Illusions Varies as a Function of the Autism-Spectrum Quotient but not in Ways Predicted by Local–Global Biases. J Autism Dev Disord. 2016;46: 2224–2239. doi:10.1007/s10803-016-2753-1

64. Shah P, Catmur C, Bird G. Emotional decision-making in autism spectrum disorder: the roles of interoception and alexithymia. Mol Autism. 2016;7: 43. doi:10.1186/s13229-016-0104-x

65. Haffey A, Press C, O’Connell G, Chakrabarti B. Autistic Traits Modulate Mimicry of Social but not Nonsocial Rewards: Autistic traits modulate mimicry of social rewards. Autism Res. 2013;6: 614–620. doi:10.1002/aur.1323

66. Ioannidis JPA. Why Most Discovered True Associations Are Inflated. Epidemiology. 2008;19: 640. doi:10.1097/EDE.0b013e31818131e7

67. Austin EJ. Personality correlates of the broader autism phenotype as assessed by the Autism Spectrum Quotient (AQ). Personal Individ Differ. 2005;38: 451–460. doi:10.1016/j.paid.2004.04.022

68. Hoekstra RA, Bartels M, Cath DC, Boomsma DI. Factor Structure, Reliability and Criterion Validity of the Autism-Spectrum Quotient (AQ): A Study in Dutch Population and Patient Groups. J Autism Dev Disord. 2008;38: 1555–1566. doi:10.1007/s10803-008-0538-x

69. Murray AL, Booth T, McKenzie K, Kuenssberg R. What range of trait levels can the Autism-Spectrum Quotient (AQ) measure reliably? An item response theory analysis. Psychol Assess. 2016;28: 673–683. doi:10.1037/pas0000215

70. Brainard DH. The Psychophysics Toolbox. Spat Vis. 1997;10: 433–436.

71. Kleiner M, Brainard D, Pelli D. What’s new in Psychtoolbox-3? Perception 36 ECVP Abstract Supplement. 2007;

72. Pelli DG. The VideoToolbox software for visual psychophysics: Transforming numbers into movies. Spat Vis. 1997;10: 437–442.

73. R Core Team. R: A Language and Environment for Statistical Computing [Internet]. Vienna, Austria: R Foundation for Statistical Computing; 2015. Available: http://www.R-project.org/

74. Singmann H, Bolker B, Westfall J, Aust F. afex: Analysis of Factorial Experiments [Internet]. 2018. Available: https://CRAN.R-project.org/package=afex

75. Halekoh U, Højsgaard S. A Kenward-Roger Approximation and Parametric Bootstrap Methods for Tests in Linear Mixed Models - The R Package pbkrtest. J Stat Softw. 2014;59. doi:10.18637/jss.v059.i09

76. Kenward MG, Roger JH. Small Sample Inference for Fixed Effects from Restricted Maximum Likelihood. Biometrics. 1997;53: 983–997. doi:10.2307/2533558

77. Bates D, Kliegl R, Vasishth S, Baayen H. Parsimonious Mixed Models. ArXiv150604967 Stat. 2015; Available: http://arxiv.org/abs/1506.04967

78. Lenth R. emmeans: Estimated Marginal Means, aka Least-Squares Means [Internet]. 2018. Available: https://CRAN.R-project.org/package=emmeans

79. Genz A, Bretz F, Hochberg Y. Approximations to multivariate t integrals with application to multiple comparison procedures. Institute of Mathematical Statistics Lecture Notes - Monograph Series. Beachwood, Ohio, USA: Institute of Mathematical Statistics; 2004. pp. 24–32. doi:10.1214/lnms/1196285623

80. Hothorn T, Bretz F, Westfall P. Simultaneous Inference in General Parametric Models. Biom J. 2008;50: 346–363. doi:10.1002/bimj.200810425

81. Jones PR, Landin L, McLean A, Juni MZ, Maloney LT, Nardini M, et al. Efficient visual information sampling develops late in childhood. J Exp Psychol Gen. 2019;148: 1138–1152. doi:10.1037/xge0000629

82. Maris E, Oostenveld R. Nonparametric statistical testing of EEG- and MEG-data. J Neurosci Methods. 2007;164: 177–190. doi:10.1016/j.jneumeth.2007.03.024

83. Bogacz R, Brown E, Moehlis J, Holmes P, Cohen JD. The physics of optimal decision making: A formal analysis of models of performance in two-alternative forced-choice tasks. Psychol Rev. 2006;113: 700–765. doi:10.1037/0033-295X.113.4.700

