## Supporting Information for "Autistic traits influence the strategic diversity of information sampling: insights from two-stage decision models"

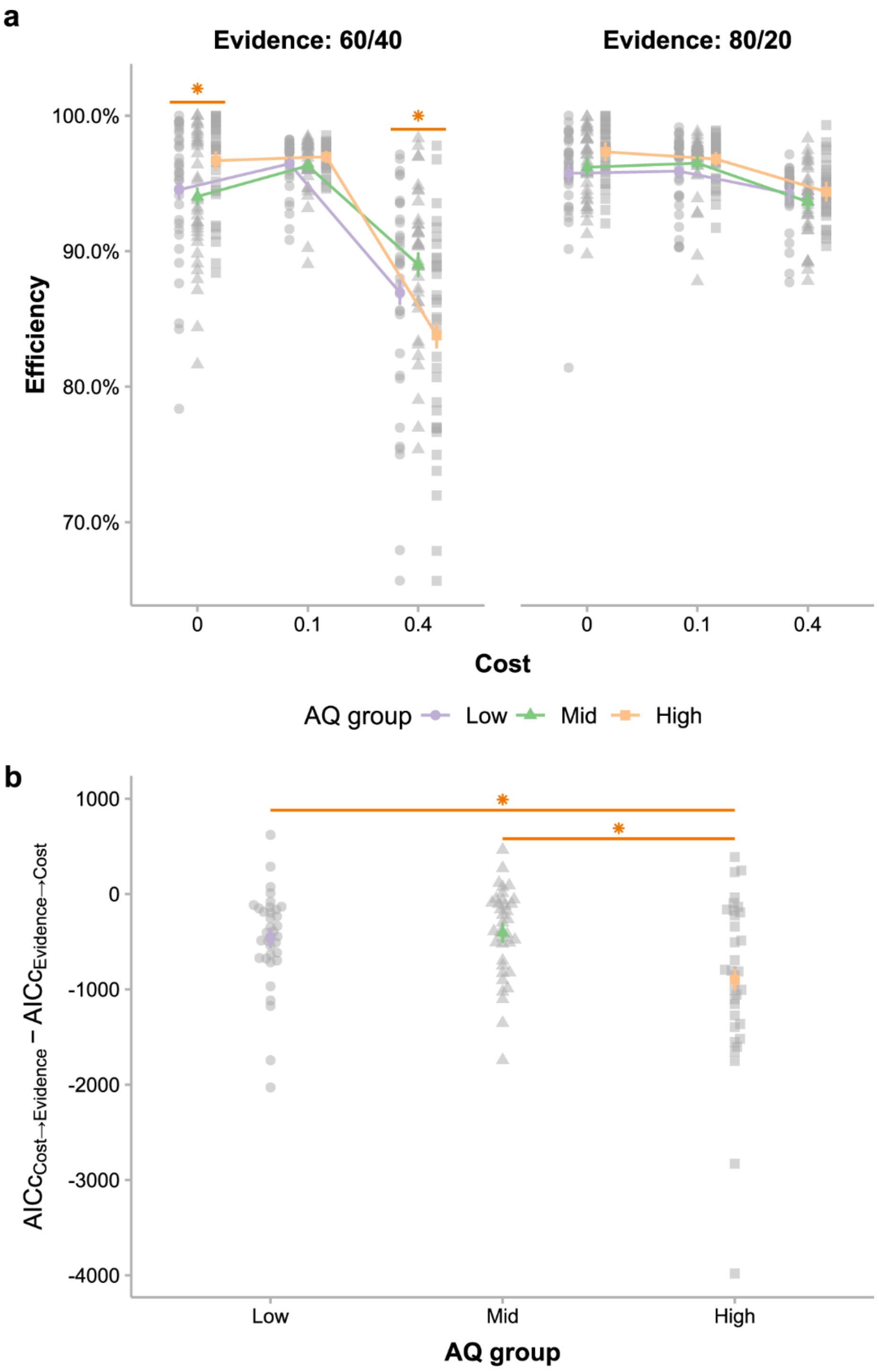


**S1 Fig. Replication of two analyses in the main text based on the tripartite division of AQ.** (a) Similar to what we have found for the linear effect of AQ on efficiency, we have found a significant three-way interaction of AQ groups, cost and evidence conditions ($F_{4,200.83}$ = 11.03, $p$ < .001). The simple main effect showed the group with high AQ had higher efficiency in the zero-cost, low-evidence condition, compared to the other two groups with low and middle AQ scores ($F_{2,132.62}$ = 4.27, $p$ = .016; *post hoc* comparison: Low – Mid: $t_{132.62}$ = 0.53, $p$ = .86, Low –

High: $t_{132.62}$ = -2.23, $p$ = .07, Mid – High: $t_{132.62}$ = -2.77, $p$ = .018, $p$s were corrected by single-step adjustment, see Methods). Meanwhile, the group with high AQ had significantly lower efficiency in the high-cost, low-evidence condition (simple main effect: $F_{2,117.67}$ = 8.18, $p$ < .001; *post hoc* comparison: Low – Mid: $t_{118.45}$ = -1.60, $p$ = .25, Low – High: $t_{117.29}$ = 2.42, $p$ = .044, Mid – High: $t_{117.25}$ = 4.02, $p$ < .001). All these results were consistent with those reported in the main text based on regressions (see Fig 2b). (b) Participants with different levels of autistic traits significantly differed in $AICc_{\text{cost}\rightarrow\text{evidence}} - AICc_{\text{evidence}\rightarrow\text{cost}}$ ($F_{2,101}$=5.96, $p$ = .004), with the value of the high-AQ group smaller than those of the low-AQ group ($t_{101}$ = -2.81, $p$ = .017) and the middle-AQ group ($t_{101}$=-3.175, $p$ = .006). This is consistent with the negative correlation between AQ and $AICc_{\text{cost}\rightarrow\text{evidence}} - AICc_{\text{evidence}\rightarrow\text{cost}}$ (see Fig 6a). In both (a) and (b), colored lines represent group means and semi-transparent gray symbols represent individual participants. Different shapes of symbols are for different AQ groups: circles for low-AQ, triangles for middle-AQ, and squares for high-AQ. Error bars denote model-based standard errors. Dark orange asterisks and lines indicate significant simple main effects ($p$ < .05).

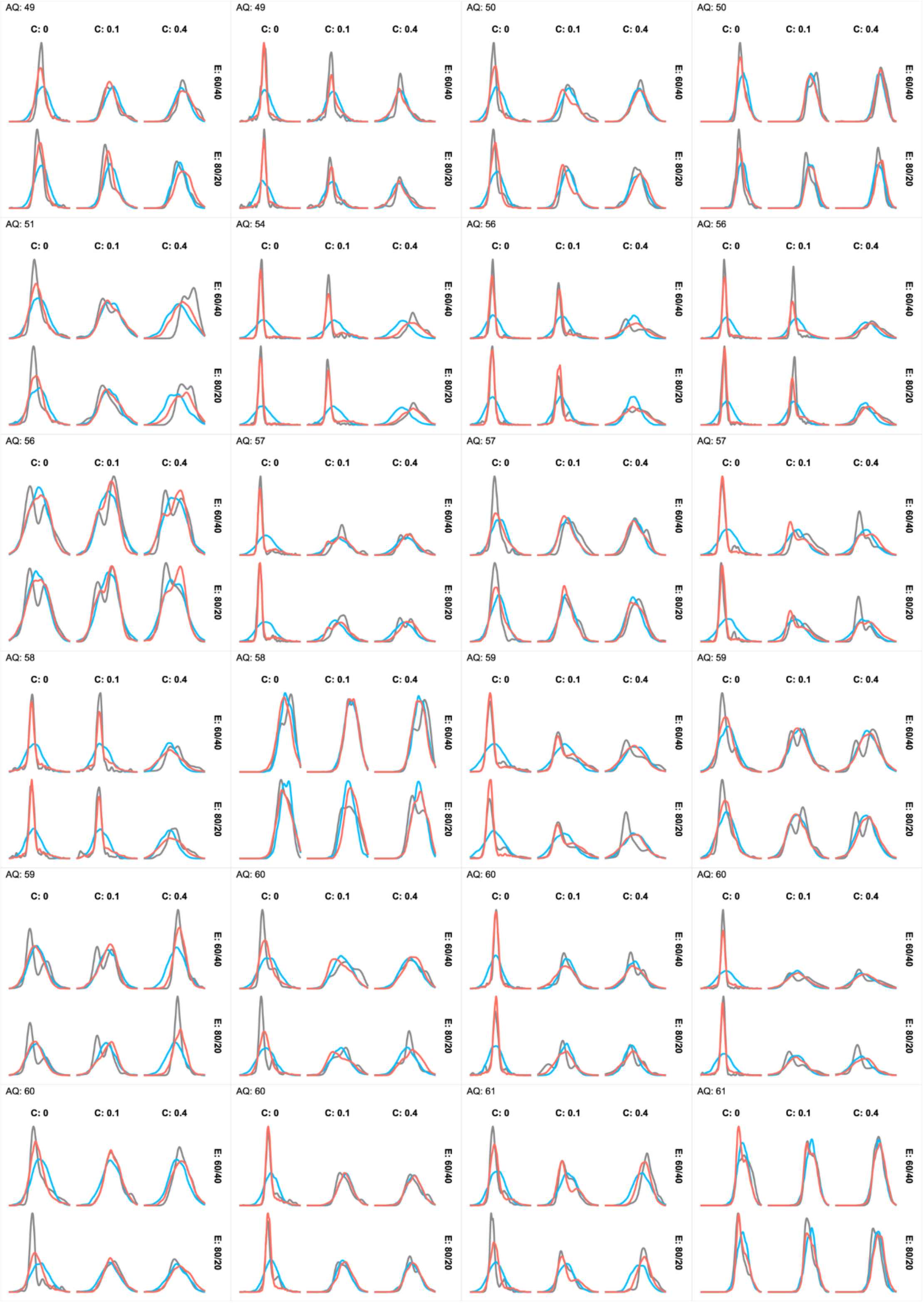
AQ: 49
C: 0
C: 0.1
C: 0.4
E: 60/40
E: 80/20
AQ: 49
C: 0
C: 0.1
C: 0.4
E: 60/40
E: 80/20
AQ: 50
C: 0
C: 0.1
C: 0.4
E: 60/40
E: 80/20
AQ: 50
C: 0
C: 0.1
C: 0.4
E: 60/40
E: 80/20
AQ: 51
C: 0
C: 0.1
C: 0.4
E: 60/40
E: 80/20
AQ: 54
C: 0
C: 0.1
C: 0.4
E: 60/40
E: 80/20
AQ: 56
C: 0
C: 0.1
C: 0.4
E: 60/40
E: 80/20
AQ: 56
C: 0
C: 0.1
C: 0.4
E: 60/40
E: 80/20
AQ: 56
C: 0
C: 0.1
C: 0.4
E: 60/40
E: 80/20
AQ: 57
C: 0
C: 0.1
C: 0.4
E: 60/40
E: 80/20
AQ: 57
C: 0
C: 0.1
C: 0.4
E: 60/40
E: 80/20
AQ: 57
C: 0
C: 0.1
C: 0.4
E: 60/40
E: 80/20
AQ: 58
C: 0
C: 0.1
C: 0.4
E: 60/40
E: 80/20
AQ: 58
C: 0
C: 0.1
C: 0.4
E: 60/40
E: 80/20
AQ: 59
C: 0
C: 0.1
C: 0.4
E: 60/40
E: 80/20
AQ: 59
C: 0
C: 0.1
C: 0.4
E: 60/40
E: 80/20
AQ: 59
C: 0
C: 0.1
C: 0.4
E: 60/40
E: 80/20
AQ: 60
C: 0
C: 0.1
C: 0.4
E: 60/40
E: 80/20
AQ: 60
C: 0
C: 0.1
C: 0.4
E: 60/40
E: 80/20
AQ: 60
C: 0
C: 0.1
C: 0.4
E: 60/40
E: 80/20
AQ: 60
C: 0
C: 0.1
C: 0.4
E: 60/40
E: 80/20
AQ: 60
C: 0
C: 0.1
C: 0.4
E: 60/40
E: 80/20
AQ: 61
C: 0
C: 0.1
C: 0.4
E: 60/40
E: 80/20
AQ: 61
C: 0
C: 0.1
C: 0.4
E: 60/40
E: 80/20

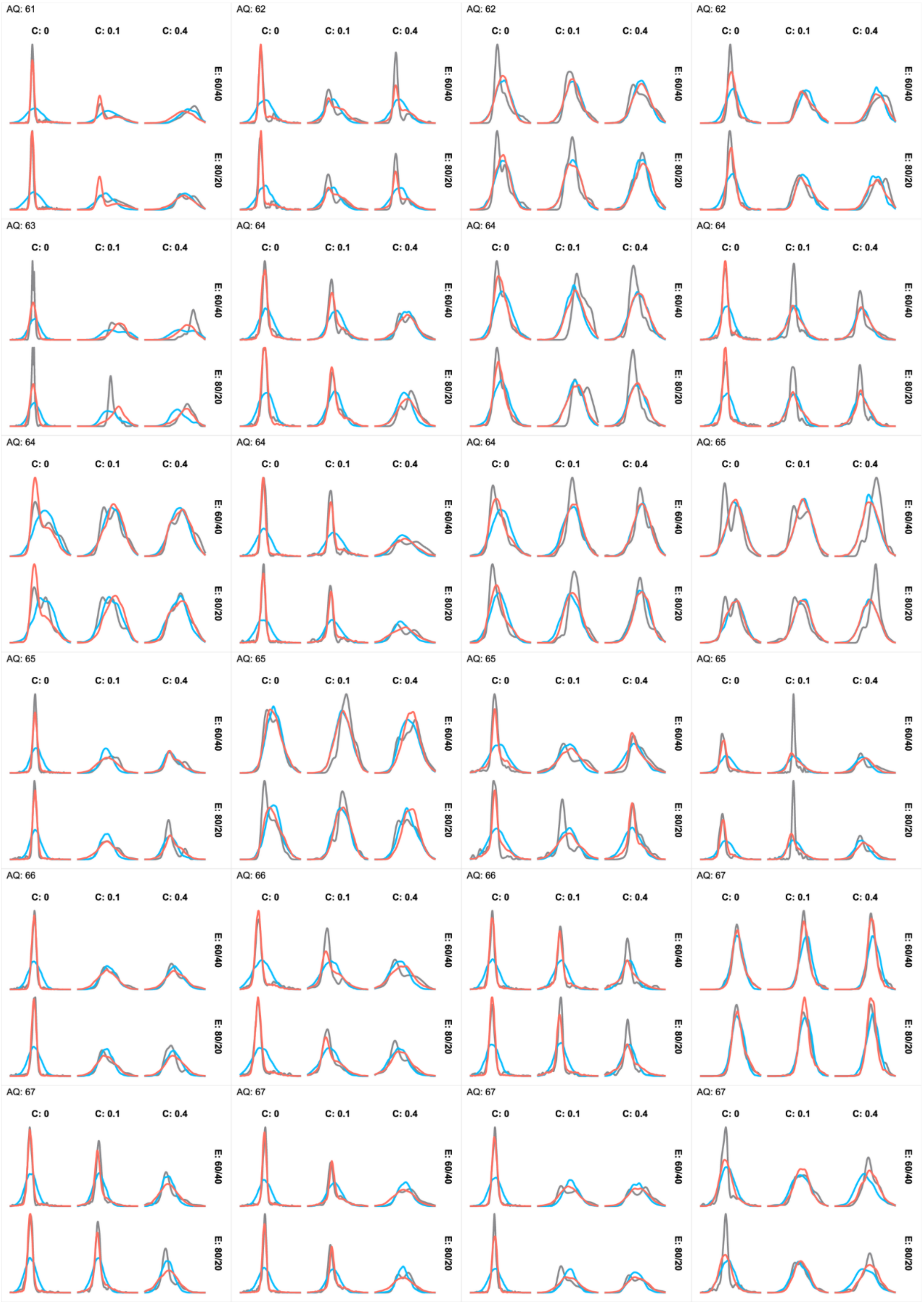
AQ: 61
C: 0
C: 0.1
C: 0.4
E: 60/40
E: 80/20
AQ: 62
C: 0
C: 0.1
C: 0.4
E: 60/40
E: 80/20
AQ: 62
C: 0
C: 0.1
C: 0.4
E: 60/40
E: 80/20
AQ: 62
C: 0
C: 0.1
C: 0.4
E: 60/40
E: 80/20
AQ: 63
C: 0
C: 0.1
C: 0.4
E: 60/40
E: 80/20
AQ: 64
C: 0
C: 0.1
C: 0.4
E: 60/40
E: 80/20
AQ: 64
C: 0
C: 0.1
C: 0.4
E: 60/40
E: 80/20
AQ: 64
C: 0
C: 0.1
C: 0.4
E: 60/40
E: 80/20
AQ: 64
C: 0
C: 0.1
C: 0.4
E: 60/40
E: 80/20
AQ: 64
C: 0
C: 0.1
C: 0.4
E: 60/40
E: 80/20
AQ: 64
C: 0
C: 0.1
C: 0.4
E: 60/40
E: 80/20
AQ: 65
C: 0
C: 0.1
C: 0.4
E: 60/40
E: 80/20
AQ: 65
C: 0
C: 0.1
C: 0.4
E: 60/40
E: 80/20
AQ: 65
C: 0
C: 0.1
C: 0.4
E: 60/40
E: 80/20
AQ: 65
C: 0
C: 0.1
C: 0.4
E: 60/40
E: 80/20
AQ: 65
C: 0
C: 0.1
C: 0.4
E: 60/40
E: 80/20
AQ: 66
C: 0
C: 0.1
C: 0.4
E: 60/40
E: 80/20
AQ: 66
C: 0
C: 0.1
C: 0.4
E: 60/40
E: 80/20
AQ: 66
C: 0
C: 0.1
C: 0.4
E: 60/40
E: 80/20
AQ: 67
C: 0
C: 0.1
C: 0.4
E: 60/40
E: 80/20
AQ: 67
C: 0
C: 0.1
C: 0.4
E: 60/40
E: 80/20
AQ: 67
C: 0
C: 0.1
C: 0.4
E: 60/40
E: 80/20
AQ: 67
C: 0
C: 0.1
C: 0.4
E: 60/40
E: 80/20
AQ: 67
C: 0
C: 0.1
C: 0.4
E: 60/40
E: 80/20

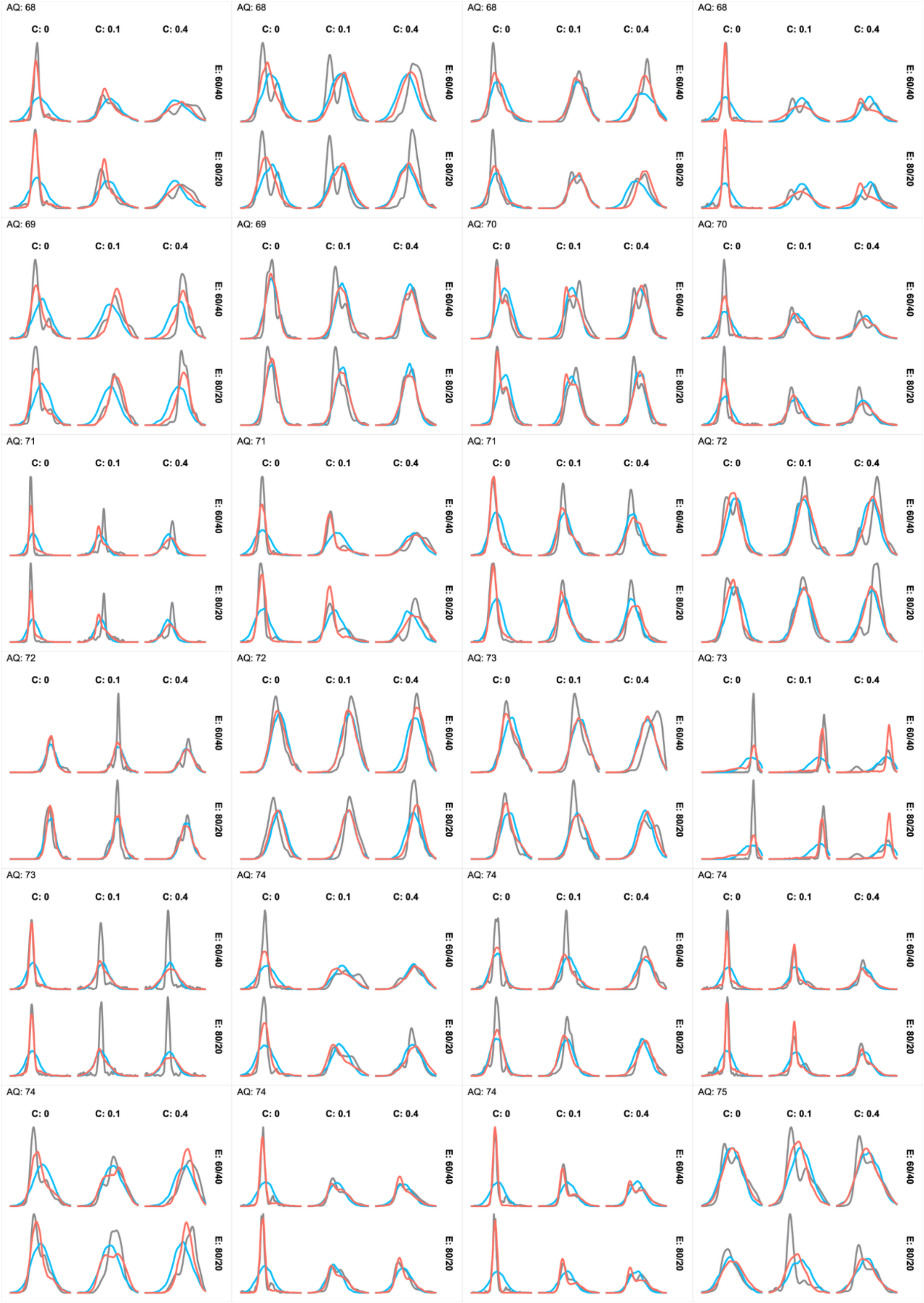
AQ: 68
C: 0
C: 0.1
C: 0.4
E: 60/40
E: 80/20
AQ: 68
AQ: 68
AQ: 68
AQ: 69
AQ: 69
AQ: 70
AQ: 70
AQ: 71
AQ: 71
AQ: 71
AQ: 72
AQ: 72
AQ: 72
AQ: 73
AQ: 73
AQ: 73
AQ: 74
AQ: 74
AQ: 74
AQ: 74
AQ: 74
AQ: 74
AQ: 75

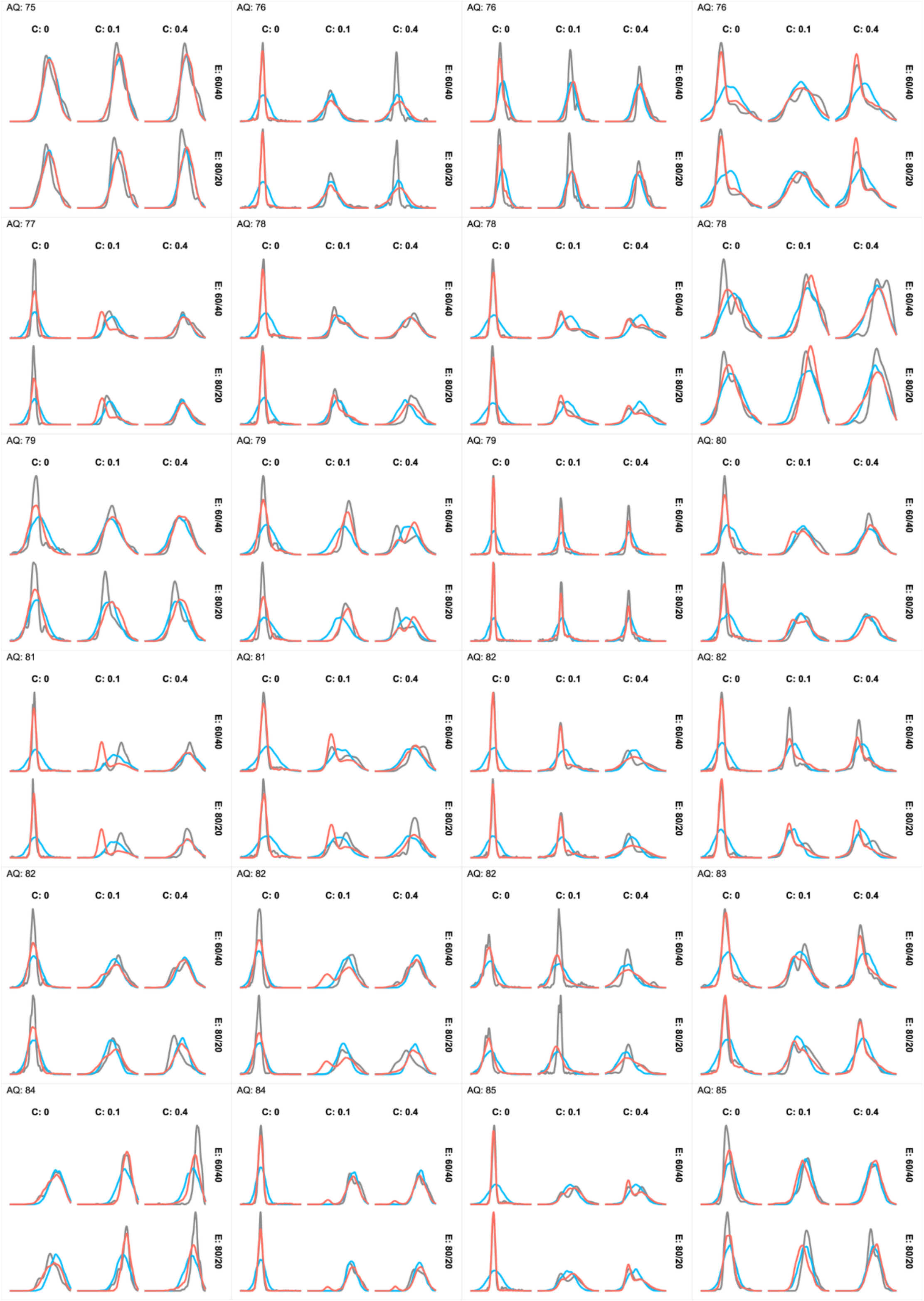

AQ: 75
AQ: 76
AQ: 76
AQ: 76
AQ: 77
AQ: 78
AQ: 78
AQ: 78
AQ: 79
AQ: 79
AQ: 79
AQ: 80
AQ: 81
AQ: 81
AQ: 82
AQ: 82
AQ: 82
AQ: 82
AQ: 82
AQ: 83
AQ: 84
AQ: 84
AQ: 85
AQ: 85
C: 0
C: 0.1
C: 0.4
E: 60/40
E: 80/20

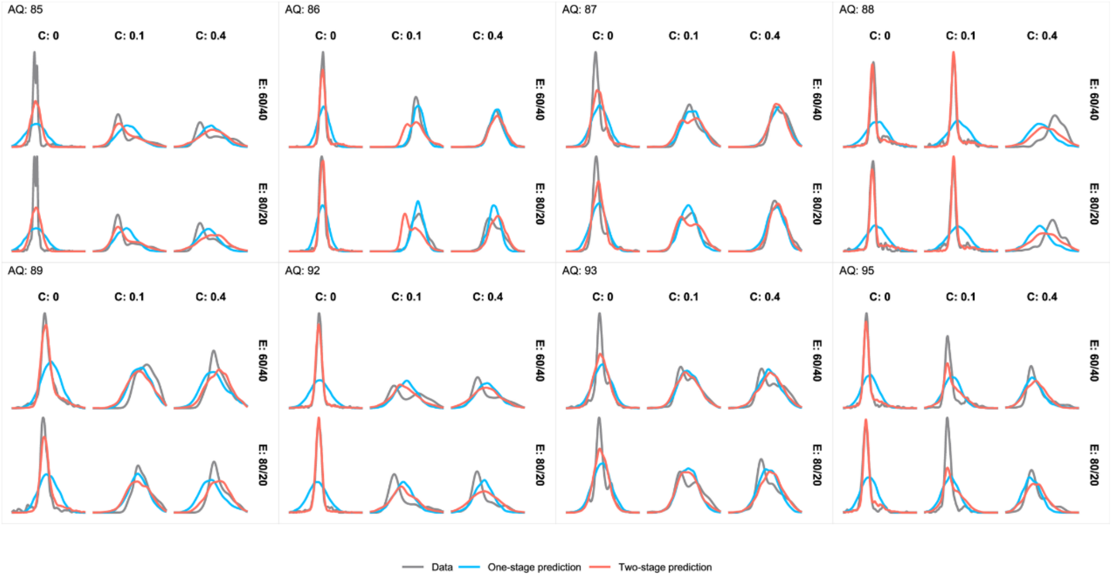


**S2 Fig. Decision time distributions of individual participants in each cost and evidence condition: data vs. model predictions (extended to 5 pages).**

Gray, blue, and red lines respectively denote data, the best one-stage model predictions, and the best two-stage model predictions. Each panel is for one participant, with each of its sub-panels for one cost and evidence condition. Panels are arranged by participants' AQ (marked at the top-left corner) ascendingly from left to right and from top to bottom. For most participants, the observed DT distributions were bimodal and were better predicted by the best-fit one-stage model than by the best-fit one-stage model. C: Cost, E: Evidence.

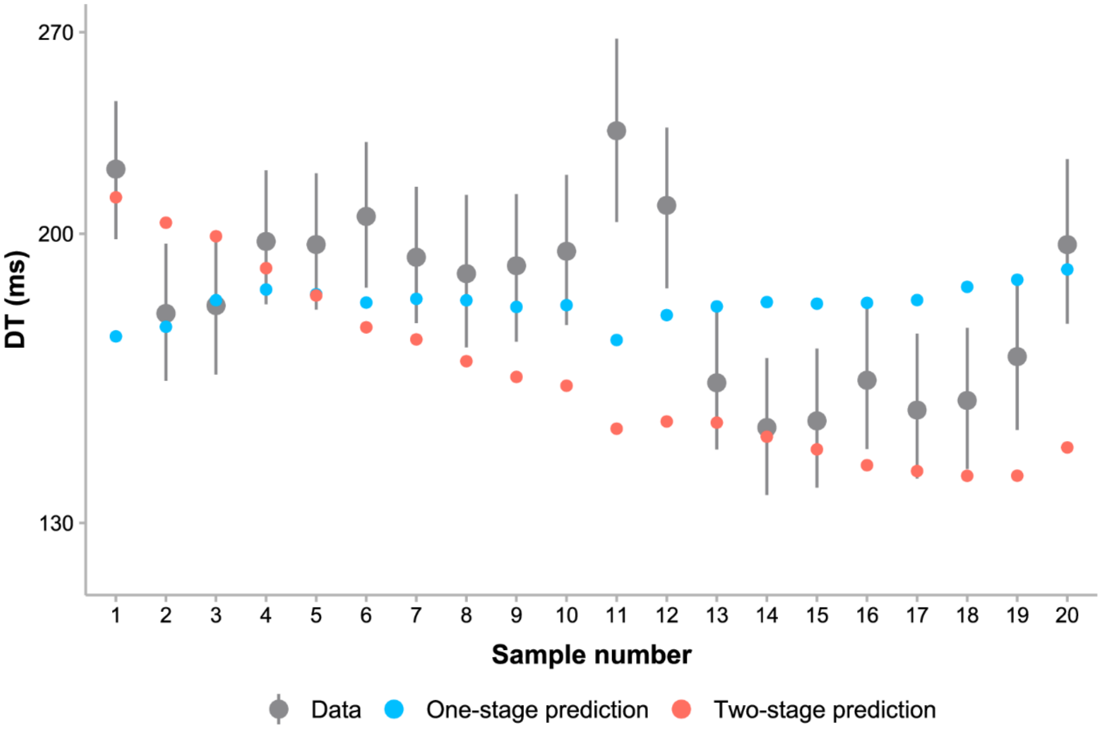


**S3 Fig. Decision time as a function of sample number: data vs. model predictions.**
The observed decision times had a significant decreasing trend with the increase of sample number ($t$ = -12.26, $p$ < .001), which was captured by the best two-stage model (red dots) but not by the best one-stage model (blue dots).

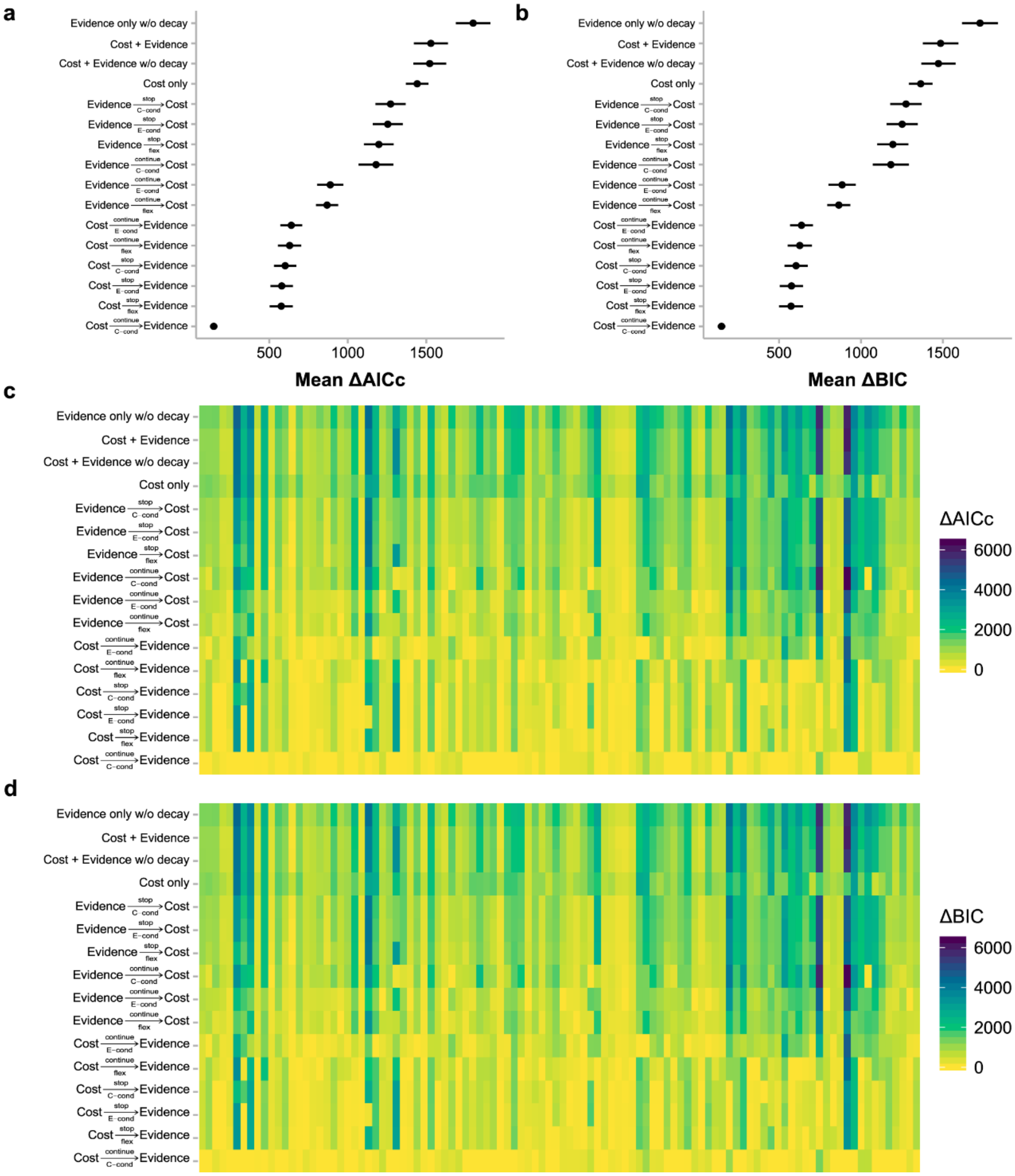


**S4 Fig. Model comparisons based on AICc and BIC and individual participants' model evidence.**

(a-b) Mean ΔAICc and ΔBIC for every model. ΔAICc or ΔBIC for a specific model was calculated for each participant with respect to the participant's best-fitting model (i.e. lowest AICc or BIC) and then averaged across participants. Error bars denote standard errors. Model comparisons based on AICc and BIC led to almost the same results. (c-d) Individual participants' ΔAICc and ΔBIC for every model. In the heatmaps, each column is for one participant, arranged in ascending order of AQ from left to right. Each row is for one model, arranged in the same order as in a-b.

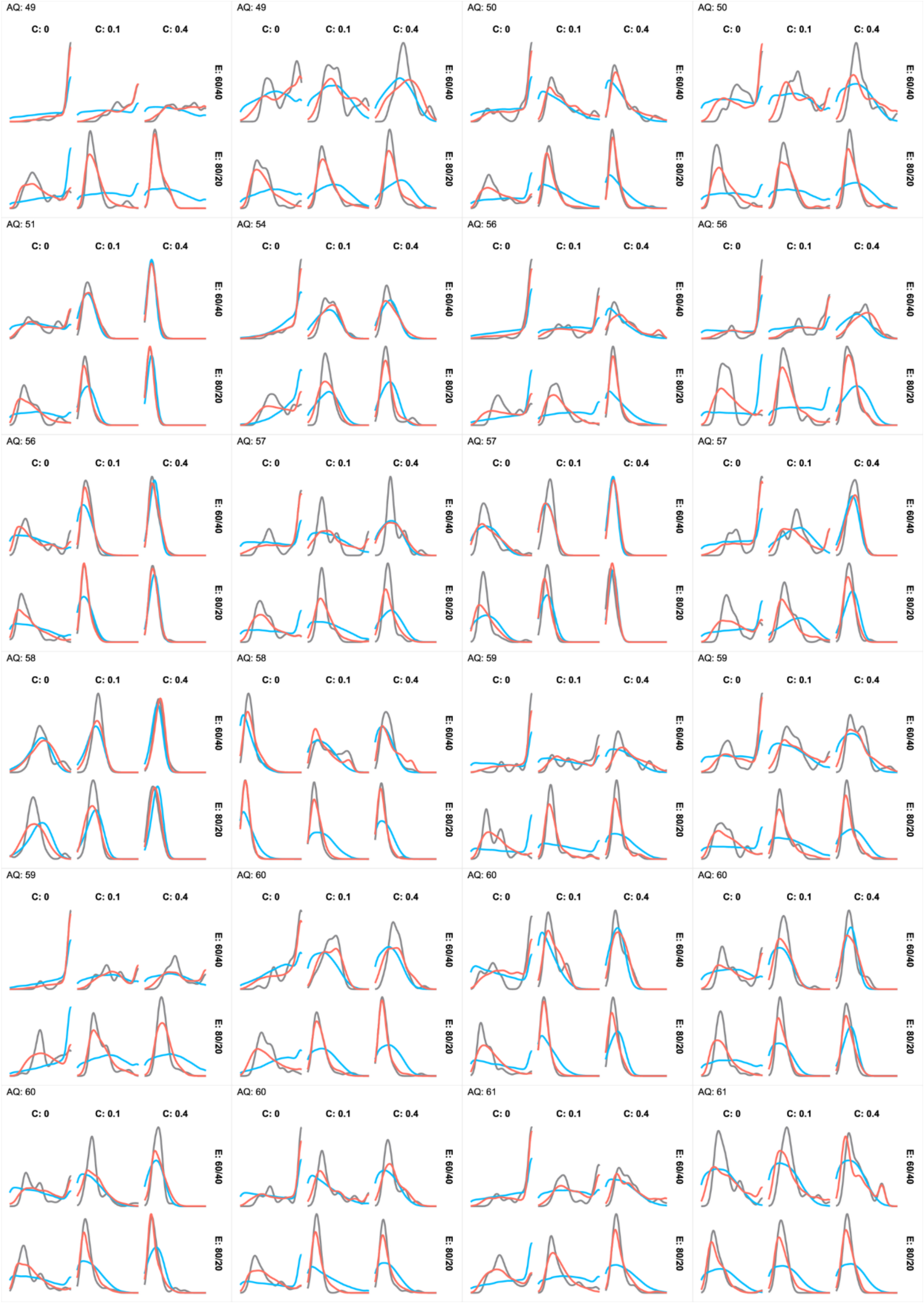
AQ: 49
AQ: 49
AQ: 50
AQ: 50
AQ: 51
AQ: 54
AQ: 56
AQ: 56
AQ: 56
AQ: 57
AQ: 57
AQ: 57
AQ: 58
AQ: 58
AQ: 59
AQ: 59
AQ: 59
AQ: 60
AQ: 60
AQ: 60
AQ: 60
AQ: 60
AQ: 61
AQ: 61
C: 0
C: 0.1
C: 0.4
E: 60/40
E: 80/20

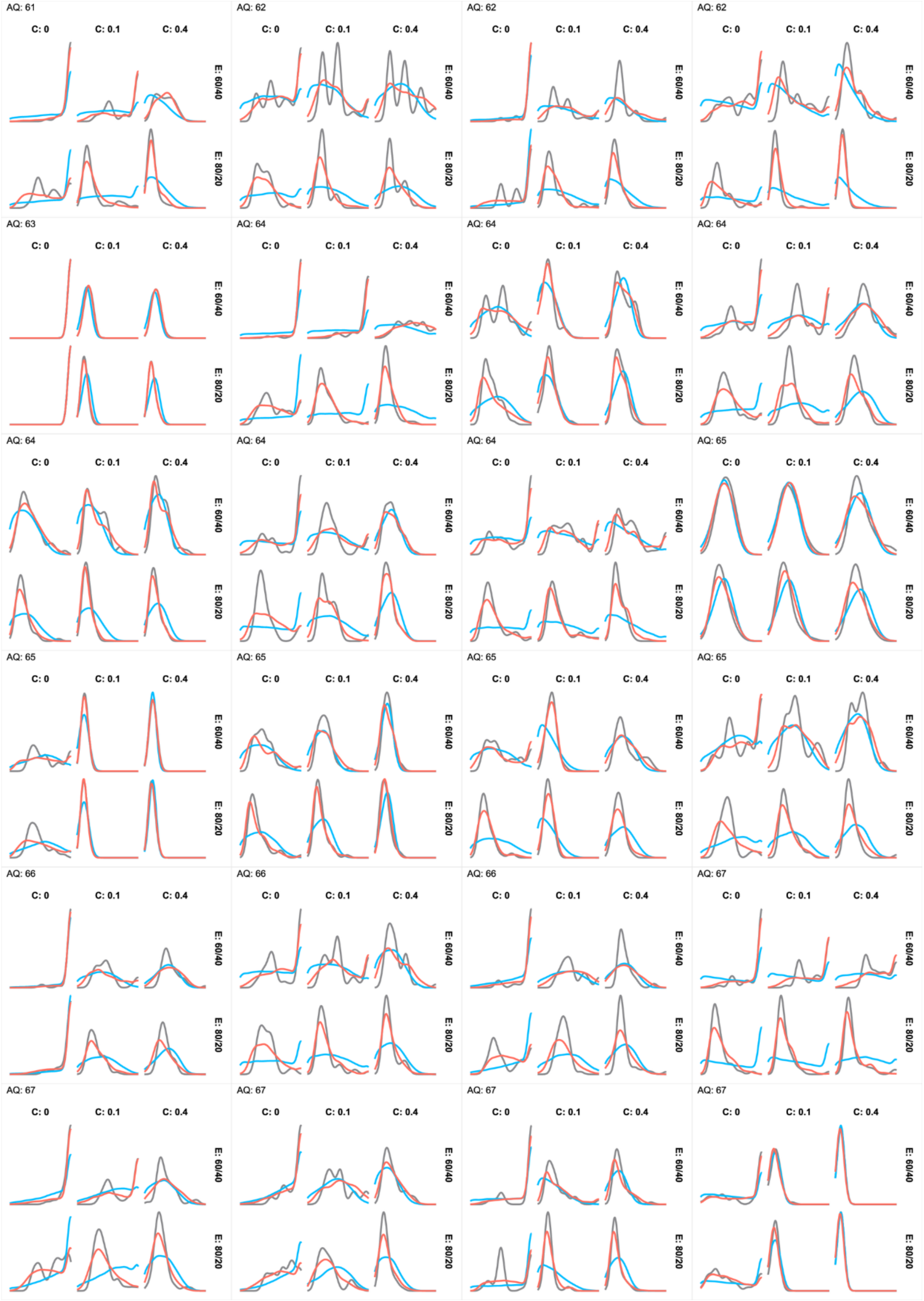
AQ: 61
AQ: 62
AQ: 62
AQ: 62
AQ: 63
AQ: 64
AQ: 64
AQ: 64
AQ: 64
AQ: 64
AQ: 64
AQ: 65
AQ: 65
AQ: 65
AQ: 65
AQ: 65
AQ: 66
AQ: 66
AQ: 66
AQ: 67
AQ: 67
AQ: 67
AQ: 67
AQ: 67
C: 0
C: 0.1
C: 0.4
E: 60/40
E: 80/20

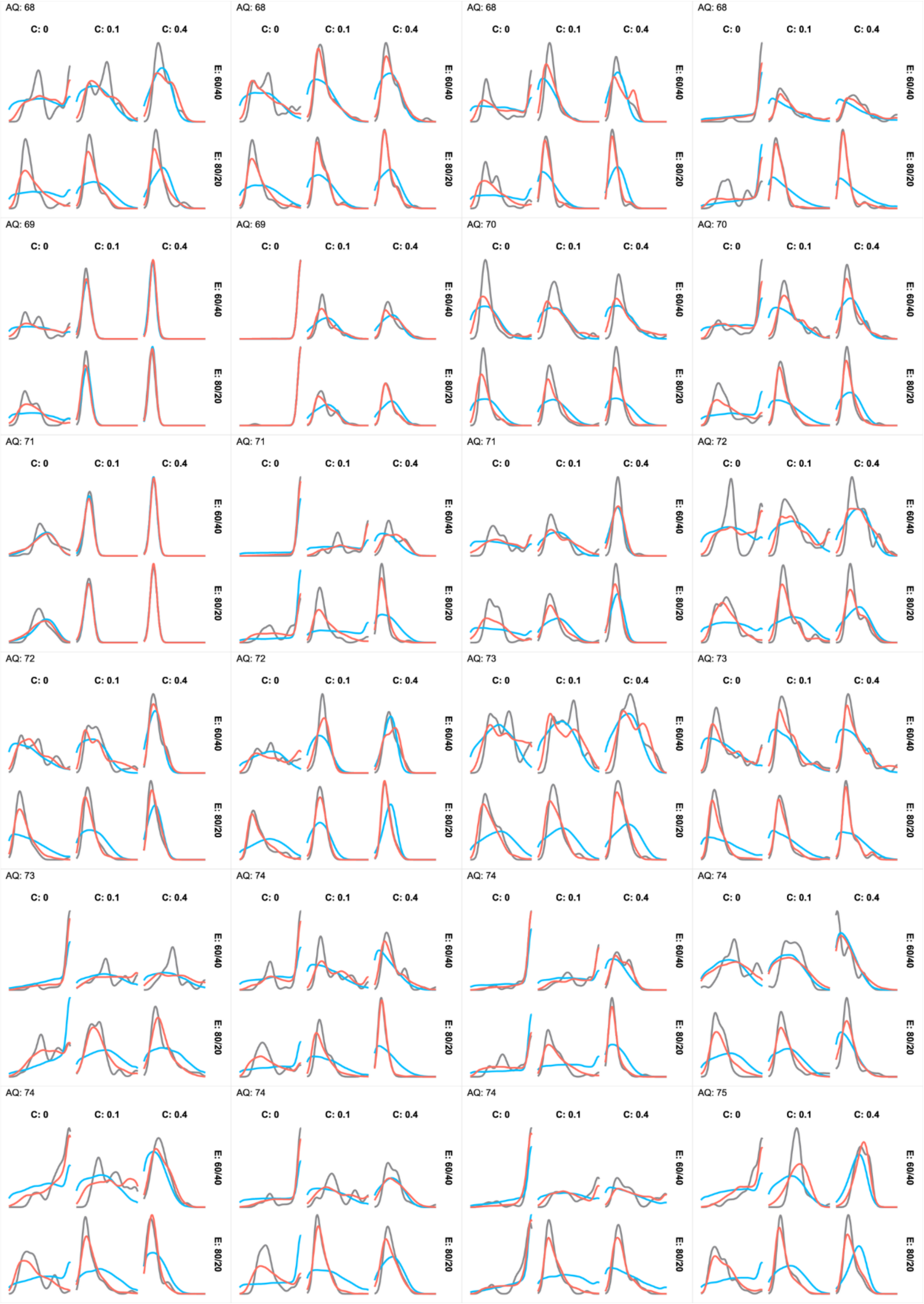

AQ: 68
C: 0
C: 0.1
C: 0.4
E: 60/40
E: 80/20
AQ: 68
C: 0
C: 0.1
C: 0.4
E: 60/40
E: 80/20
AQ: 68
C: 0
C: 0.1
C: 0.4
E: 60/40
E: 80/20
AQ: 68
C: 0
C: 0.1
C: 0.4
E: 60/40
E: 80/20
AQ: 69
C: 0
C: 0.1
C: 0.4
E: 60/40
E: 80/20
AQ: 69
C: 0
C: 0.1
C: 0.4
E: 60/40
E: 80/20
AQ: 70
C: 0
C: 0.1
C: 0.4
E: 60/40
E: 80/20
AQ: 70
C: 0
C: 0.1
C: 0.4
E: 60/40
E: 80/20
AQ: 71
C: 0
C: 0.1
C: 0.4
E: 60/40
E: 80/20
AQ: 71
C: 0
C: 0.1
C: 0.4
E: 60/40
E: 80/20
AQ: 71
C: 0
C: 0.1
C: 0.4
E: 60/40
E: 80/20
AQ: 72
C: 0
C: 0.1
C: 0.4
E: 60/40
E: 80/20
AQ: 72
C: 0
C: 0.1
C: 0.4
E: 60/40
E: 80/20
AQ: 72
C: 0
C: 0.1
C: 0.4
E: 60/40
E: 80/20
AQ: 73
C: 0
C: 0.1
C: 0.4
E: 60/40
E: 80/20
AQ: 73
C: 0
C: 0.1
C: 0.4
E: 60/40
E: 80/20
AQ: 73
C: 0
C: 0.1
C: 0.4
E: 60/40
E: 80/20
AQ: 74
C: 0
C: 0.1
C: 0.4
E: 60/40
E: 80/20
AQ: 74
C: 0
C: 0.1
C: 0.4
E: 60/40
E: 80/20
AQ: 74
C: 0
C: 0.1
C: 0.4
E: 60/40
E: 80/20
AQ: 74
C: 0
C: 0.1
C: 0.4
E: 60/40
E: 80/20
AQ: 74
C: 0
C: 0.1
C: 0.4
E: 60/40
E: 80/20
AQ: 74
C: 0
C: 0.1
C: 0.4
E: 60/40
E: 80/20
AQ: 75
C: 0
C: 0.1
C: 0.4
E: 60/40
E: 80/20

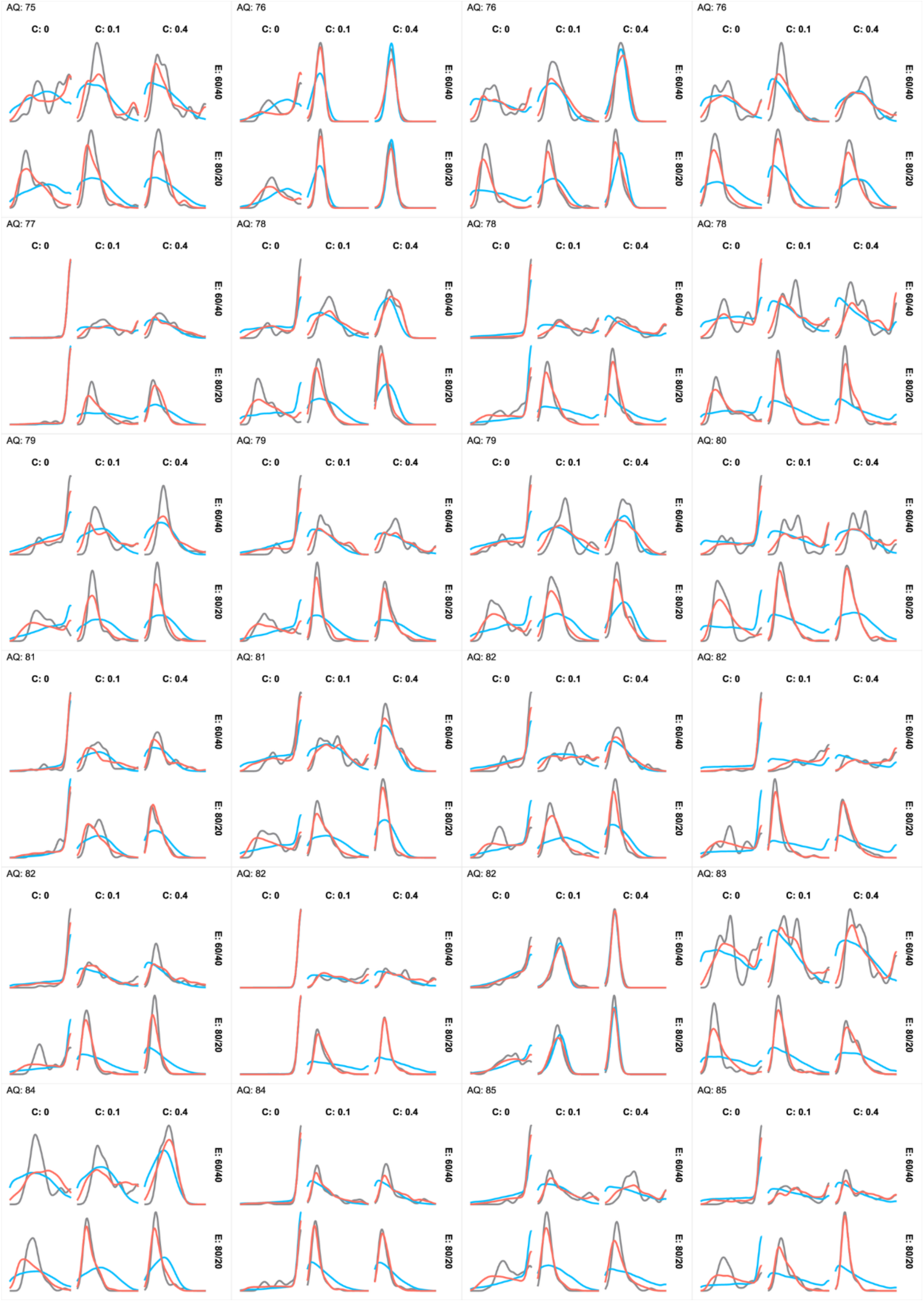
AQ: 75
C: 0
C: 0.1
C: 0.4
E: 60/40
E: 80/20
AQ: 76
C: 0
C: 0.1
C: 0.4
E: 60/40
E: 80/20
AQ: 76
C: 0
C: 0.1
C: 0.4
E: 60/40
E: 80/20
AQ: 76
C: 0
C: 0.1
C: 0.4
E: 60/40
E: 80/20
AQ: 77
C: 0
C: 0.1
C: 0.4
E: 60/40
E: 80/20
AQ: 78
C: 0
C: 0.1
C: 0.4
E: 60/40
E: 80/20
AQ: 78
C: 0
C: 0.1
C: 0.4
E: 60/40
E: 80/20
AQ: 78
C: 0
C: 0.1
C: 0.4
E: 60/40
E: 80/20
AQ: 79
C: 0
C: 0.1
C: 0.4
E: 60/40
E: 80/20
AQ: 79
C: 0
C: 0.1
C: 0.4
E: 60/40
E: 80/20
AQ: 79
C: 0
C: 0.1
C: 0.4
E: 60/40
E: 80/20
AQ: 80
C: 0
C: 0.1
C: 0.4
E: 60/40
E: 80/20
AQ: 81
C: 0
C: 0.1
C: 0.4
E: 60/40
E: 80/20
AQ: 81
C: 0
C: 0.1
C: 0.4
E: 60/40
E: 80/20
AQ: 82
C: 0
C: 0.1
C: 0.4
E: 60/40
E: 80/20
AQ: 82
C: 0
C: 0.1
C: 0.4
E: 60/40
E: 80/20
AQ: 82
C: 0
C: 0.1
C: 0.4
E: 60/40
E: 80/20
AQ: 82
C: 0
C: 0.1
C: 0.4
E: 60/40
E: 80/20
AQ: 82
C: 0
C: 0.1
C: 0.4
E: 60/40
E: 80/20
AQ: 83
C: 0
C: 0.1
C: 0.4
E: 60/40
E: 80/20
AQ: 84
C: 0
C: 0.1
C: 0.4
E: 60/40
E: 80/20
AQ: 84
C: 0
C: 0.1
C: 0.4
E: 60/40
E: 80/20
AQ: 85
C: 0
C: 0.1
C: 0.4
E: 60/40
E: 80/20
AQ: 85
C: 0
C: 0.1
C: 0.4
E: 60/40
E: 80/20

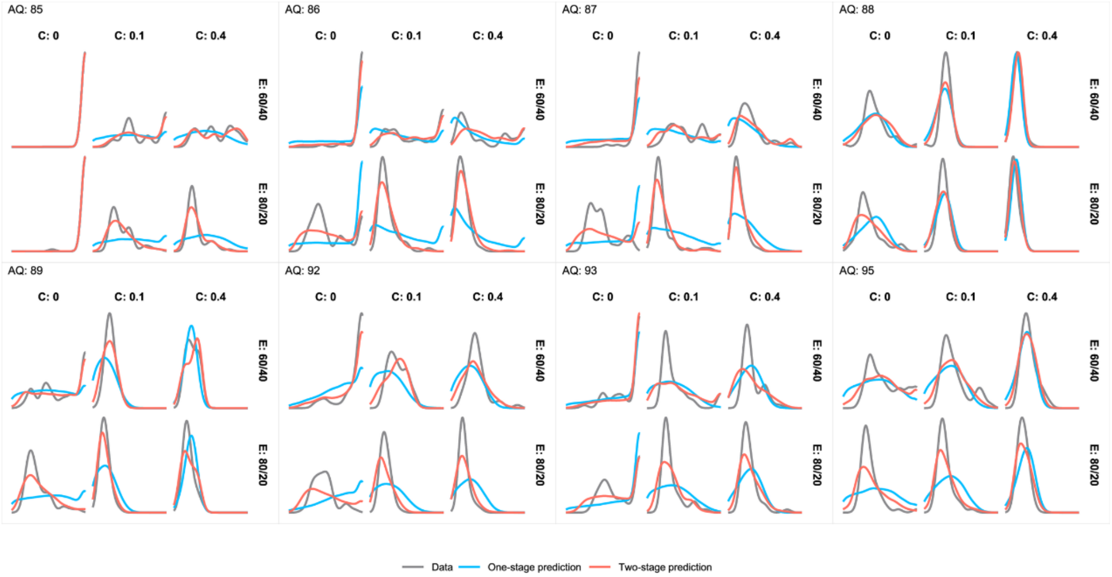


**S5 Fig. Sample size distributions of individual participants in each cost and evidence condition: data vs. model predictions (extended to 5 pages).**

Gray, blue, and red lines respectively denote data, the best one-stage model predictions, and the best two-stage model predictions. Each panel is for one participant, with each of its sub-panels for one cost and evidence condition. Panels are arranged by participants' AQ (marked at the top-left corner) ascendingly from left to right and from top to bottom. For most participants, the observed sample size distributions were better predicted by the best-fit one-stage model than by the best-fit one-stage model. C: Cost, E: Evidence.

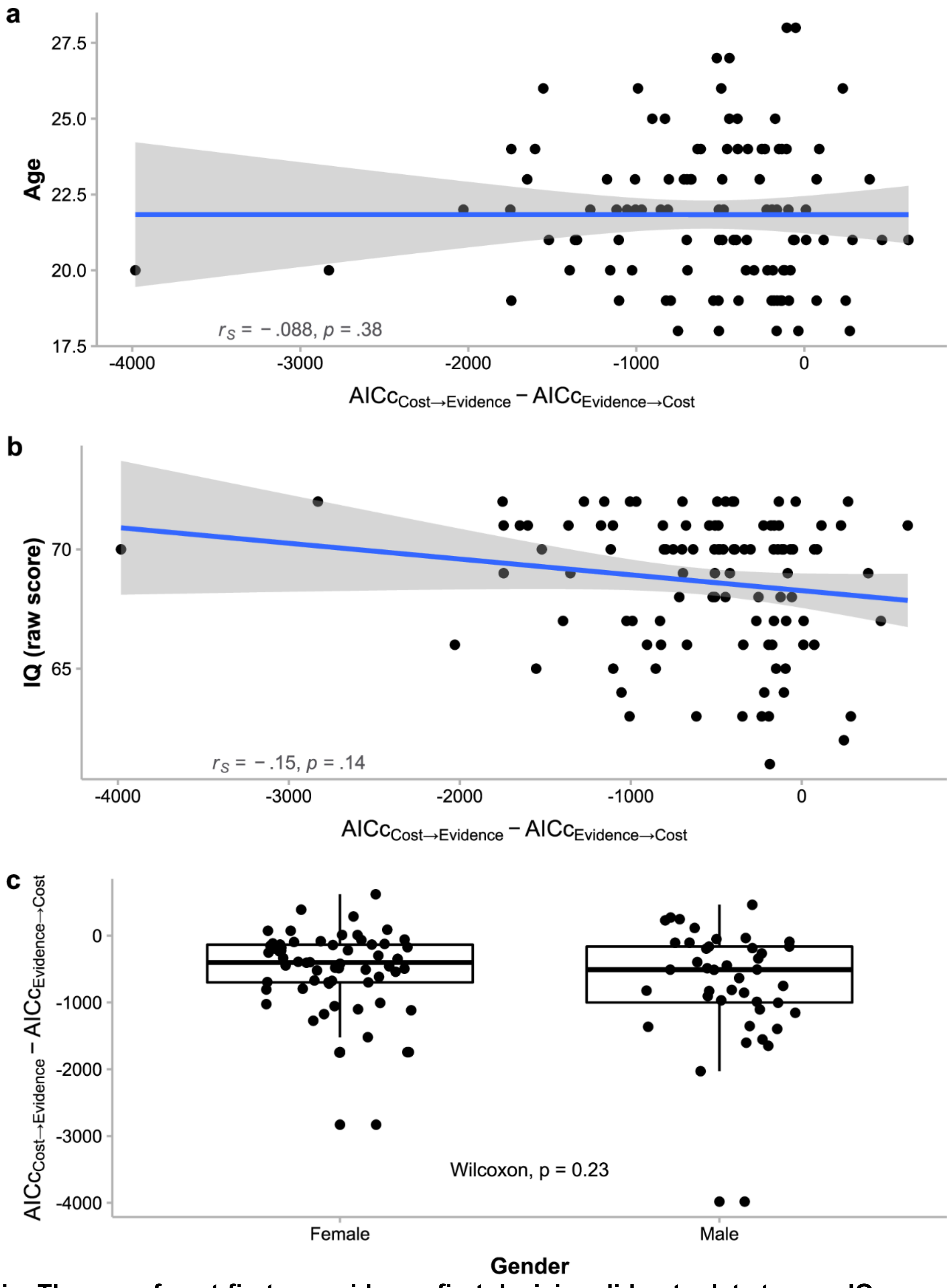


**S6 Fig. The use of cost-first vs. evidence-first decision did not relate to age, IQ, or gender.**

There were little correlations between $AICc_{\text{cost}\rightarrow\text{evidence}} - AICc_{\text{evidence}\rightarrow\text{cost}}$ and participants' age (a) or IQ score (b); $AICc_{\text{cost}\rightarrow\text{evidence}} - AICc_{\text{evidence}\rightarrow\text{cost}}$ did not differ between genders either (c).

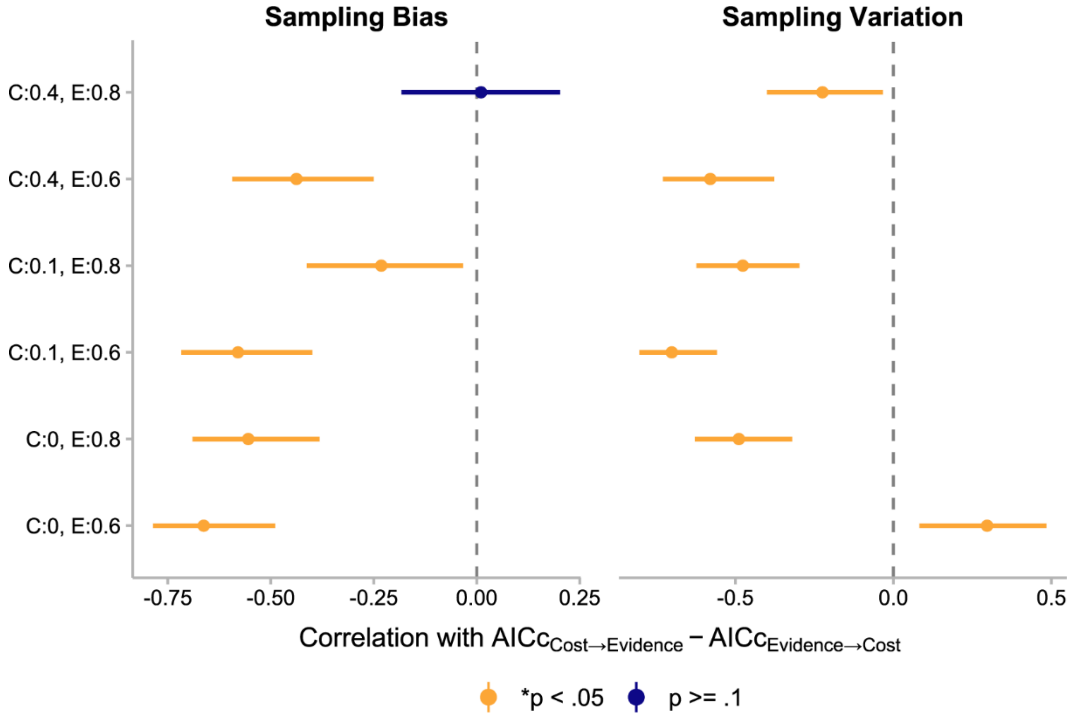


**S7 Fig. How the use of cost-first vs. evidence-first decision processes related to sampling bias and sampling variation.**

Correlations between $AICc_{\text{cost}\rightarrow\text{evidence}} - AICc_{\text{evidence}\rightarrow\text{cost}}$ and sampling bias (signed deviation from the optimal number of sampling, denoted $\overline{n_s - n_{opt}}$ ) or sampling variation (standard deviation of actual number of sampling across trials, denoted $SD(n_s)$) under different cost and evidence conditions were consistent with what we would expect if AQ affects these measures through cost-first vs. evidence-first preference in decision process. See Fig 6b for the corresponding plot for efficiency. Error bars represent 95% confidence intervals (FDR corrected). C:0 = zero-cost, C:0.1 = low-cost, C:0.4 = high-cost, E:0.6 = low-evidence, E:0.8 = high-evidence.

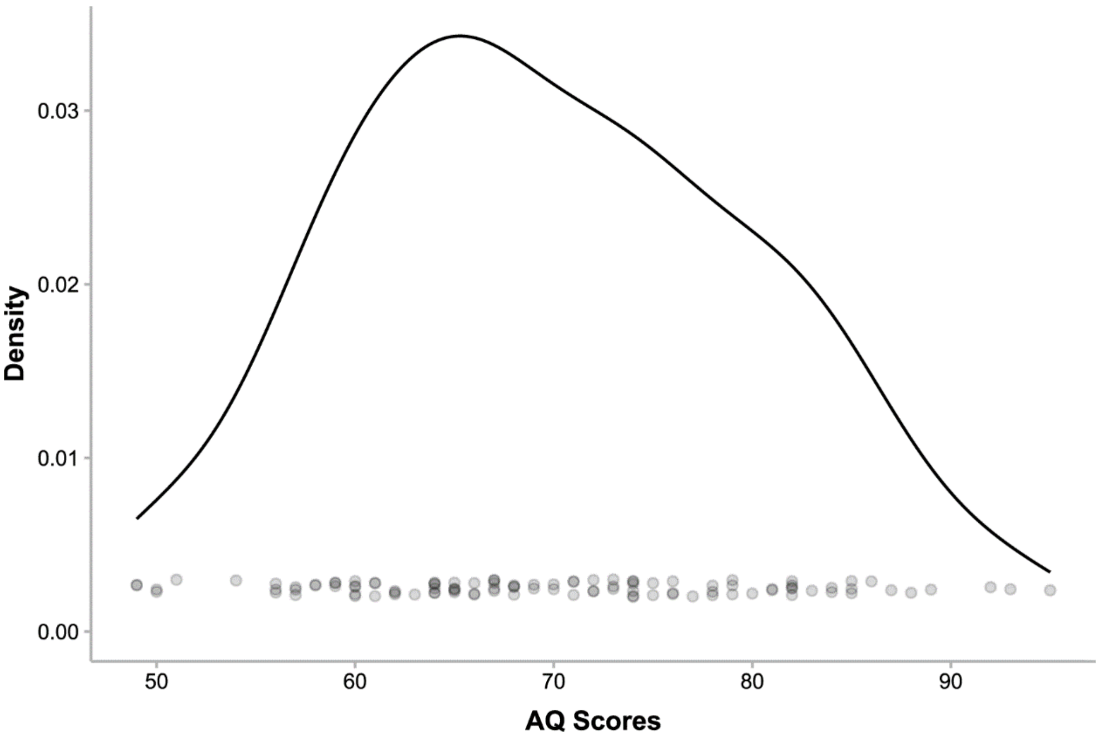


**S8 Fig. Distribution of AQ scores among the 104 participants.** Each circle denotes one participant.

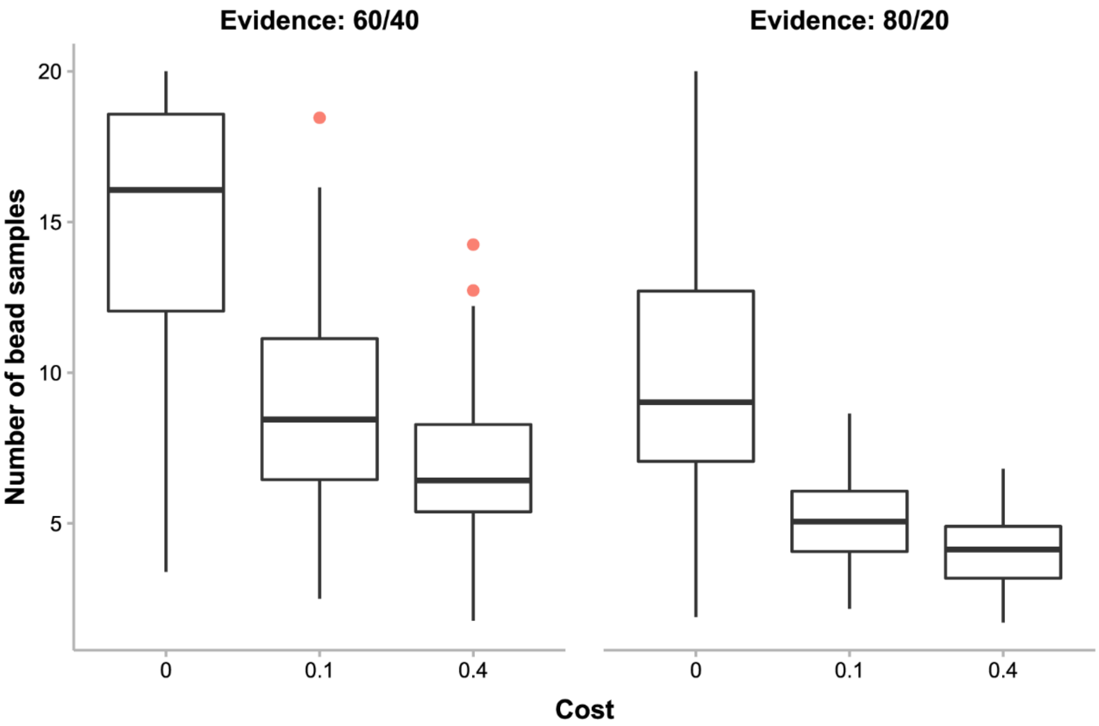


**S9 Fig. Noncompliant observations.**

Following Jones et al. [81], we identified three observations (red dots) as “likely noncompliant” in the number of bead samples for each condition based on nonparametric boxplot statistics, that is, those whose values were lower than the 1st quartile or higher than the 3rd quartile of all the observations in the condition by more than 1.5 times of the interquartile range. These observations (not participants per se) were excluded from linear mixed model analyses 1-3 (LMM 1-3, see Methods).
